# Gene Editing for *ATXN3* Inactivation in Machado-Joseph disease: CRISPR-Cas9 as a Therapeutic Alternative to TALEN-Induced Toxicity

**DOI:** 10.1101/2025.02.14.637261

**Authors:** Sara Monteiro Lopes, Miguel Monteiro Lopes, Daniela Oliveira, Magda Matos Santana, Ana Rita Fernandes, Ana Vasconcelos-Ferreira, Dina Pereira, Maria Casquinha, Clévio Nóbrega, Carlos A. Matos, Neville E. Sanjana, Patrick D. Hsu, Fei Ann Ran, Lukasz Swiech, Le Cong, Feng Zhang, Rui Jorge Nobre, Luís Pereira de Almeida

## Abstract

Machado-Joseph disease (MJD) is an autosomal dominantly-inherited neurodegenerative disorder, caused by an over-repetition of the polyglutamine-codifying region in the *ATXN3* gene. Strategies based on the suppression of the deleterious gene products have demonstrated promising results in pre-clinical studies. Nonetheless, these strategies do not target the root cause of the disease. In order to prevent the downstream toxic pathways, our goal was to develop gene editing-based strategies to permanently inactivate the human *ATXN3* gene. TALENs and CRISPR-Cas9 systems were designed to target exon 2 of this gene and functional characterization was performed in a human cell line. After the demonstration of TALEN’s and CRISPR-Cas9 efficiency on gene disruption, a sequence of each system was selected for further *in vivo* experiments. Although both TALENs and CRISPR-Cas9 systems led to a drastic reduction of ATXN3 aggregates in the striatum of a lentiviral-based mouse model of MJD/SCA3, only CRISPR-Cas9 system allowed the improvement of key neuropathological markers of the disease. Importantly, the administration of the engineered system in YAC-MJD84.2/84.2 mice mediated a delay in disease progression, when compared with non-treated littermates. These data provide the first *in vivo* evidence of the efficacy of a CRISPR-Cas9-based approach to permanently inactivate the *ATXN3* gene in the brain of two mouse models of the disease, supporting its potential as a new therapeutic avenue in the context of MJD/SCA3.

## INTRODUCTION

Machado-Joseph disease (MJD), or spinocerebellar ataxia type 3 (SCA3) ^1,2^, is one of the nine known polyglutamine (polyQ) expansion diseases. Despite rare, MJD/SCA3 is the most common autosomal dominantly-inherited ataxia worldwide ^3,4^. The disease mutation has been mapped to chromosome 14 (14q32.12) and it consists of an unstable CAG repeat expansion in the coding region of the ataxin-3 (*ATXN3*) gene, which translates into a pathogenic polyQ repeat expansion at the C-terminus of the ATXN3 protein ^5,6^. Sequences ranging from 61 to 87 have been related with the development of MJD/SCA3 ^7^. In disease conditions, the presence of an expanded polyQ tract induces conformational changes in ATXN3, which acquires toxic properties. As a result, some important cellular functions are compromised, culminating in neuronal dysfunction and degeneration in specific brain regions, such as the cerebellum, brainstem and striatum ^8–10^. Although some patients benefit from symptomatic therapy, no effective causative treatment able to prevent, cure or delay the progression of this disorder has been developed.

Considering the toxic gain-of-function acquired by the expanded ATXN3 species, strategies aiming at reducing *ATXN3* mRNA levels with resource to RNA interference ^11–18^ and antisense oligonucleotides ^19,20^ are developing rapidly and have proven efficient to slowdown the neurodegenerative cascade in several MJD/SCA3 models. Despite limiting the disease progression, the aforementioned strategies may only produce an incomplete and/or transient therapeutic effect in target cells or tissues ^14,19^ and therefore may fail to achieve a disease’s cure.

By acting at the DNA level, genome editing technologies, on the other hand, have been successfully used to permanently inactivate or correct disease-related genes, holding promise for the development of a definitive cure for inherited diseases, such as MJD/SCA3. One of these tools are transcription activator-like effector nucleases (TALENs), which are chimeric proteins that combine features of a DNA-binding domain with programmable specificity and a non-specific nuclease domain ^21–23^. The central DNA-binding domain originates from TALE proteins that are secreted by the bacterial plant pathogen *Xanthomonas* and that consists of a variable number of tandem repeats ^24–26^. Each repeat unit of the DNA-binding domain is typically 34 amino acid long. The sequence of these units is largely identical, with variations occurring primarily at amino acids 12 and 13 (repeat variable di-residues, RVDs), the identity of which dictates the binding specificity of each repeat to a single nucleotide at the target site (one-to-one correspondence) ^27–30^. The C-terminal catalytic domain consists of the FokI endonuclease, naturally found in *Flavobacterium okeanokoites,* which confers a non-specific endonuclease activity upon dimerization. Therefore, a functional pair of TALENs, spaced by a few nucleotides (14 to 20 bases), must be used in combination to allow the dimerization of FokI catalytic domains that function as molecular scissors ^31,32^.

The most recently developed gene editing platform is based on the bacterial clustered regularly interspaced short palindromic repeat (CRISPR) system that derives from *Streptococcus pyogenes.* Briefly, the bacterial CRISPR-associated (Cas) nuclease 9 (*Sp*Cas9) is recruited to the region of interest by a short single guide RNA (sgRNA) molecule that binds to a complementary 20 base pair (bp) DNA sequence, via Watson-Crick pairing. A critical feature of this system is the existence of an obligatory protospacer adjacent motif (PAM) sequence immediately downstream of the DNA target site. *Sp*Cas9 targets DNA sites flanked by 5’-NGG PAM sequences, catalyzing a double-strand break (DSB) at approximately 3 bp upstream of the PAM, through the activation of the HNH and RuvC nuclease domains ^33–36^.

Upon the introduction of double-strand breaks, gene editing is achieved through the activation of endogenous cellular repair machinery. In eukaryotic cells there are two major repair mechanisms to restore DSBs ^37^, although the error-prone non-homologous end-joining (NHEJ) constitutes the most active repair mechanism throughout the cell cycle ^37,38^. NHEJ repairs the lesion by re-joining the two cleaved ends in a process that does not require a repair template. Consequently, this repair process results in nucleotide insertions or deletions (indels) at the lesion site ^39^. When introduced into a coding sequence of a gene, these indels will often result in frameshift mutations that create premature stop codons, leading to mRNA degradation by nonsense-mediated decay or to the production of non-functional proteins ^40^. Thus, similarly to RNA silencing methods, NHEJ can be used to suppress gene function ^41–43^, with the major advantage of being a permanent approach.

In view of the urgency to develop new therapeutic strategies to counteract MJD/SCA3’s pathogenesis, we designed TALEN and CRISPR-Cas9 systems to permanently inactivate the human *ATXN3* gene. Although both DNA targeting nucleases are efficient to disrupt the *ATXN3* gene, a marked amelioration of the neuropathological signs was observed in a lentiviral-based mouse model of MJD/SCA3 only when using the CRISPR-Cas9 system. The therapeutic potential of the developed CRISPR-Cas9 strategy was further tested in a transgenic mouse model of MJD/SCA3, through the injection of adeno-associated virus (AAV) encoding the therapeutic CRISPR-Cas9 system directly into the cerebrospinal fluid. Notably, a halt in the progression of MJD/SCA3 was observed in treated mice, suggesting that this CRISPR-Cas9-based tool may constitute a potential therapeutic strategy for this incurable disorder.

## RESULTS

### Engineered TALEN pairs effectively target the human *ATXN3* gene, reducing protein expression in a human cell line

To permanently block the human *ATXN3* gene expression, we used a TALEN-based strategy to introduce a DSB in exon 2 of the gene. Since TALENs’ catalytic domain is only active as a dimer ^32^, three functional pairs of properly oriented TALENs (pair A/D, B/E and C/F) were designed taking into account the following criteria: i) all of the selected TALEN-binding sites must begin with a thymine ^29,30^; ii) each TALEN must target a 20 bp DNA sequence ^23^, and iii) left and right TALEN target sites must be spaced by approximately 14-20 bases to facilitate Fokl dimerization. Importantly, the RVD modules used in our constructs for DNA recognition (NI, HD, NG and NH) are predicted to display high binding specificities, which minimizes mismatch tolerability. Moreover, the use of obligatory heterodimeric FokI domains (ELD:KKR mutants) has also been shown to improve on-target specificity ^44,45^. Accordingly, the designed TALEN pairs target a total length of 55 bp (TALEN A/D), 58 bp (TALEN B/E) and 59 bp (TALEN C/F) in the *ATXN3* gene, being spaced by 15 bp, 18 bp and 19 bp, respectively. Relative positions and target sequences can be found in Figure 1A and 1B, respectively.

**Figure 1.**
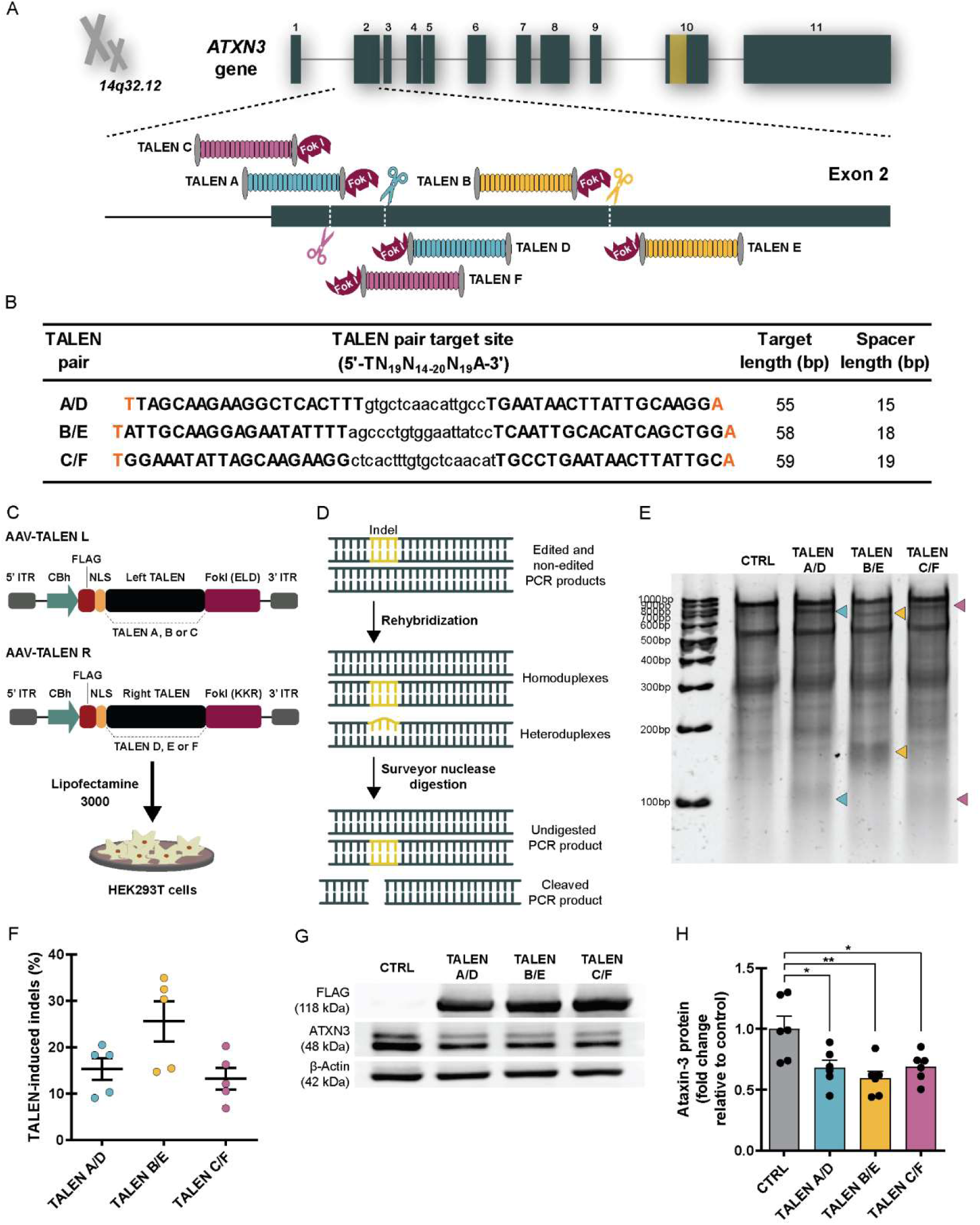
Customized TALEN pairs efficiently target the human *ATXN3* gene, leading to a consequent reduction in the expression of the resultant protein in a human cell line. **(A)** Schematic representation of TALEN target sites in the exon 2 of the human *ATXN3* gene (relative positions represented). Each left and right TALEN recognize the sense and antisense DNA strands, respectively, being separated by a spacer sequence of variable length (14 to 20 bp). Between the pair of binding sites, FokI catalytic domains dimerize and cut the DNA (colored scissors), stimulating genome editing via NHEJ repair pathway for the permanent inactivation of *ATXN3* gene expression. Colors indicate each of the three TALEN pairs (TALEN A/D; TALEN B/E; TALEN C/F). **(B)** Each pair of TALENs target a 5’-TN_19_N_14-20_N_19_A-3’ region (N = A, G, T or C), where left TALEN targets 5’-TN_19_-3’ (sequence specified in upper case letters) and right TALEN targets the antisense strand of 5’-N_19_A-3’ (sequence specified in upper case letters), being spaced by 14 to 20 bases (sequence specified in lower case letters). **(C)** For the functional validation of the generated constructs, HEK293T cells were co-transfected with each pair of TALENs (left and right), using Lipofectamine 3000. Cells were harvested 48 hours after transfection. Cells transfected with the transfection reagent alone (mock transfection) were used as negative controls. **(D)** Schematic representation of the Surveyor nuclease assay. After the PCR amplification of the region surrounding TALEN target site, PCR product includes a mixture of both TALEN-modified (containing indels) and unmodified genomic DNA regions. These PCR products are melted and slowly reannealed, creating a mixture of homo- and heteroduplexes (imperfect annealing). The Surveyor nuclease detects mismatches in the imperfectly annealed heteroduplexes, promoting the enzymatic cleavage of the DNA. Cleavage fragments can be then analyzed by gel electrophoresis. **(E)** PAGE gel showing the result of Surveyor nuclease assay used to determine TALENs’ cleavage efficiency in HEK293T cells. Lane 1: DNA ladder (GeneRuler 100 bp); Lane 2: mock-transfected cells; Lanes 3-5: Cells co-transfected with TALEN A/D, TALEN B/E and TALEN C/F, respectively. Arrowheads indicate the expected Surveyor-cleaved DNA products. **(F)** The estimated indel occurrence within the human *ATXN3* locus is calculated based on the fraction of Surveyor-cleaved DNA (represented in percentage, n=5). **(G-H)** Western blot analysis reveals a significant decrease in the ATXN3 protein levels after TALEN targeting in comparison with the experimental control (n=6). Results are presented as fold change relative to mock-transfected cells. Optical densitometry analysis of ATXN3 fractions were normalized with β-actin signals. Statistical significance was evaluated with one-way ANOVA with Dunnett’s post hoc test (*p<0.05 and **p<0.01). Data are expressed as mean ± SEM. **Abbreviations**: **ITR** (invert terminal repeat); **CBh** (chicken beta-actin promoter); **FLAG** (FLAG octapeptide tag); **NLS** (nuclear localization signal); **TALEN** (transcription activator-like effector nuclease); **FokI** (catalytic domain from the endonuclease isolated from *Flavobacterium okeanokoites*); **FokI-ELD** (FokI domain with mutations Q486E, I499L and N496D); **FokI-KKR** (FokI domain with mutations E490K, I538K and H537R).

After the hierarchical ligation of each TALEN, the generated constructs were sequence-verified for the correct monomer assembly (Supplemental Figure S1). The efficiency of each pair of TALENs to edit the *ATXN3* gene was evaluated by transfecting HEK293T cells with the generated TALEN-codifying plasmids (Figure 1C) and using the Surveyor nuclease assay ^46,47^ (Figure 1D). PAGE gel electrophoresis of Surveyor digested products confirmed the ability of the TALEN pairs to modify the *ATXN3* gene in this cell line (Figure 1E). TALEN’s cleavage efficiency (% of indels) was calculated based on the fraction of cleaved DNA (Figure 1F, TALEN A/D: 15.28 ± 2.33%; TALEN B/E: 25.57 ± 4.36%; TALEN C/F: 13.18 ± 2.32%). As a consequence, a significant reduction in the levels of endogenous ATXN3 protein was observed in the western blot analysis (Figure 1G and H, CTRL: 1.00 ± 0.10; TALEN A/D: 0.68 ± 0.06; TALEN B/E: 0.59 ± 0.06; TALEN C/F: 0.69 ± 0.05), confirming TALENs’ gene disrupting efficiency.

Considering that TALEN pair B and E achieved the greatest reduction in ATXN3 protein levels in this human cell line, this TALEN pair was selected for subsequent *in vivo* experiments.

### TALENs designed to permanently inactivate the human *ATXN3* gene induce a toxic response in the mouse brain

Viral vectors are particularly suitable options for the delivery of site-specific nuclease-encoded genes ^48–50^ and have been described as efficient tools for *in vivo* gene delivery in CNS disorders ^51^. Mosaic recombinant adeno-associated virus co-expressing capsid proteins from AAV2 and AAV1 (rAAV1/2) were used to deliver intact TALEN genes *in vivo.* To address their limited cargo capacity (∼4.5 kb) ^52^, two rAAV vectors were used to deliver each of the two TALEN arms (TALEN B and TALEN E) in equal amounts into the striatum of adult mice (Supplemental Figure S2A). The detection of FLAG tag-positive cells in the brain parenchyma confirmed the efficient viral transduction of the injected striata 4 weeks after rAAV delivery (Supplemental Figure S2B).

To assess the efficiency of TALENs to mediate gene editing in the adult brain, we took advantage of our previously developed lentiviral-based mouse model of MJD/SCA3 ^53,54^. The generation of this model involves the stereotaxic injection of lentiviral vectors encoding for the human mutant ATXN3 protein with 72 glutamines, under the control of PGK promoter (LV-PGK-*ATXN3* 72Q), in the mouse striatum. Seven week-old mice were stereotaxically co-injected with LV-PGK-*ATXN3* 72Q and rAAV1/2 vectors encoding either ssGFP (4×10^9^ vg total, left striatum) or TALEN pair B and E (4×10^9^ vg total at a 1:1 ratio, right striatum, Figure 2A-B). Four weeks after stereotaxic injection, the western blot showed a reduction in the levels of mutant ATXN3 protein species in striatal tissues transduced with TALEN B/E (Figure 2C), indicating the successful silencing of the *ATXN3* gene.

**Figure 2.**
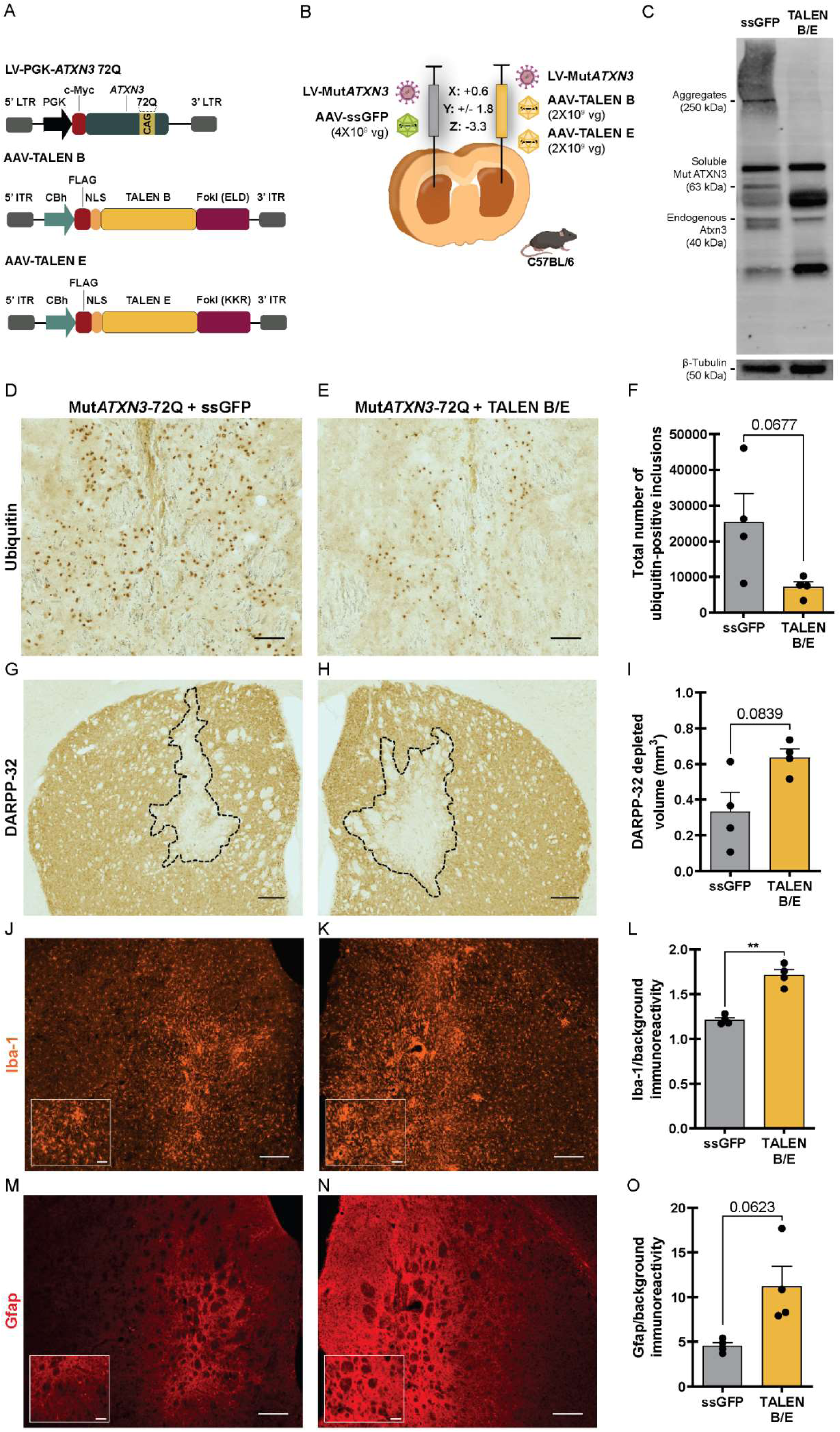
TALEN pair B and E reduces mutant ATXN3 accumulation *in vivo*, but it does not associate with striatal neuroprotection in a lentiviral-based mouse model of MJD/SCA3. **(A)** Schematic representation of the viral constructs used for the production of lentiviral and adeno-associated viral particles for *in vivo* delivery. The lentiviral construct LV-PGK-*ATXN3* 72Q drives the expression of the human mutant ATXN3 protein (Myc-tagged) with 72 glutamines, under the control of PGK promoter. The adeno-associated viral constructs drive the expression of either TALEN B (rAAV1/2-TALEN B, FLAG-tagged) or TALEN E (rAAV1/2-TALEN E, FLAG-tagged), under the control of CBh promoter. **(B)** Schematic representation of the stereotaxic injection of viral vectors in the striatum of seven-week-old C57BL/6 mice. Lentivirus encoding for the human mutant ATXN3 protein with 72Q (LV-PGK-*ATXN3* 72Q) and rAAVs encoding ssGFP were co-injected in the left hemisphere, serving as experimental control. In the contralateral hemisphere rAAVs encoding for TALEN B and TALEN E were injected, along with the human mutant ATXN3 codifying vector (LV-PGK-*ATXN3* 72Q). Four weeks after surgery animals were sacrificed. **(C)** Western blot analysis shows the reduction of ATXN3 aggregates, together with the mutant ATXN3 soluble form in TALEN-edited striata (data not quantified). **(D-E)** Immunohistochemical peroxidase staining, performed in brain slices, using anti-ubiquitin antibody. Scale bar, 50 µm. TALEN B/E injected hemispheres exhibit a tendency for a decreased number of ubiquitin-positive inclusions, when compared to the control ssGFP left hemispheres, as quantified in **(F)**. **(G-H)** Immunohistochemical analysis using anti-DARPP-32 antibody for lesion identification. Scale bar, 200 µm. Treated hemispheres, injected with the TALEN pair B and E, show a tendency for increased DARPP-32 depleted volume (mm^3^) in comparison with the contralateral hemispheres, as quantified in **(I)**. **(J-K)** Iba-1 immunoreactivity in the mouse striatum, indicates that microglial recruitment is increased in the TALEN-edited hemisphere (K), when compared with the contralateral non-edited control (J), as quantified in **(L)**. Scale bar, 200 µm in general view images and 50 µm in detail magnifications. **(M-N)** Gfap immunoreactivity in the mouse striatum display a tendency for an increased astrocytic activation in *ATXN3*-edited striatum (N) in comparison with the *ATXN3*-non-edited striatum, injected with ssGFP (M), as quantified in **(O)**. Scale bar, 200 µm in general view images and 50 µm in detail magnifications. Statistical significance was evaluated with paired Student’s t-test (**p<0.01, n=4). Data are expressed as mean ± SEM. **Abbreviations**: **LTR** (long terminal repeat); **PGK** (phosphoglycerate kinase promoter); **c-Myc** (short peptide sequence derived from the C-terminal region of the c-Myc gene); **DARPP-32** (Dopamine- and cyclic AMP-regulated phosphoprotein 32); **Iba-1** (ionized calcium binding adaptor molecule 1); **Gfap** (glial fibrillary acidic protein).

Misfolded mutant ATXN3 tends to aggregate, and the presence of neuronal intranuclear inclusions is one of the hallmarks of the disease ^55^, that is recapitulated in this mouse model. Immunohistochemical analysis of coronal sections from injected animals showed a tendency for reduction in the total number of ubiquitin-positive inclusions in hemispheres injected with TALEN B/E compared to the contralateral control hemispheres (Figure 2D-F; Mut*ATXN3*-72Q + ssGFP: 25432 ± 7845 aggregates *vs* Mut*ATXN3*-72Q + TALEN B/E: 7158 ± 1409 aggregates).

Histologically, this animal model is also characterized by the loss of DARPP-32 immunoreactivity ^53,56^. DARPP-32 is a regulator of dopamine receptor signaling ^57^ and the downregulation of its expression is indicative of early neuronal dysfunction. Unexpectedly, immunostaining for DARPP-32 revealed a tendency for an increased volume of depleted staining in TALEN B/E-injected hemispheres compared to control hemispheres (Figure 2G-I; Mut*ATXN3*-72Q + ssGFP: 0.33 ± 0.11 mm^3^ *vs* Mut*ATXN3*-72Q + TALEN B/E: 0.64 ± 0.05 mm^3^), suggesting an aggravation of neuronal dysfunction.

To investigate whether this could be due to an exacerbated inflammatory response, we performed an immunohistochemical analysis for Iba-1 and Gfap, markers of microglial and astrocytic activation, respectively. Consistent with the observation of increased neuronal dysfunction in TALEN-treated hemispheres, a local increase of Iba-1 (Figure 2K-L) and Gfap (Figure 2N-O) immunoreactivity was also detected in these hemispheres when compared with untreated controls (Figure 2J and M).

Together, these results show that the TALEN pair B and E efficiently disrupted the human mutant *ATXN3* gene *in vivo*, leading to the clearance of ubiquitin-positive inclusions. Unexpectedly, this gene editing approach did not prevent the neuropathological phenotype commonly observed in this lentiviral-based mouse model of MJD/SCA3. In fact, neuronal dysfunction and neuroinflammation seemed to be aggravated in treated hemispheres.

### CRISPR-Cas9 system targeting the human *ATXN3* gene effectively reduces ATXN3 protein levels in a human cell line

Given the lack of tolerability of the previously developed TALEN-based system, a CRISPR-Cas9-based approach was designed to edit exon 2 of the human *ATXN3* gene, and its therapeutic potential was evaluated in the context of MJD/SCA3.

Four 20-nucleotide guide sequences were designed for this purpose: sgKO.1, sgKO.2, sgKO.3 and sgKO.4 (relative positions and target sequences specified in Figure 3A). These sequences were initially cloned into a lentiviral backbone, which co-expresses both *Sp*Cas9 (FLAG-tagged) and the sgRNA scaffold (Figure 3B). To evaluate the gene editing efficiency of the generated guide sequences, HEK293T cells were transfected with the sgRNA-expressing plasmids (Figure 3B). The visualization of the Surveyor products upon electrophoretic separation demonstrated that all the designed sgRNAs were capable of inducing DSBs at the target site, effectively modifying the *ATXN3* gene in this cell line (Figure 3C). Locus modification efficiencies (percentage of indels) were determined (Figure 3D; sgKO.1: 53.38 ± 0.54%; sgKO.2: 49.33 ± 2.28%; sgKO.3: 56.26 ± 2.22%; sgKO.4: 56.04 ± 1.59%), showing the high editing efficiency of each of the sgRNAs. As a consequence of the expression of the guide sequences in HEK293T cells, a significant and robust reduction in ATXN3 protein levels was observed in CRISPR-edited cells, as determined by western blot (Figure 3E and F; sgCTRL: 1.00 ± 0.17 vs sgKO.1: 0.50 ± 0.08, sgKO.2: 0.43 ± 0.13, sgKO.3: 0.44 ± 0.10, sgKO.4: 0.48 ± 0.09). These findings suggest that the developed gene editing approach effectively inactivates the *ATXN3* gene as a result of NHEJ-mediated DSB repair.

**Figure 3.**
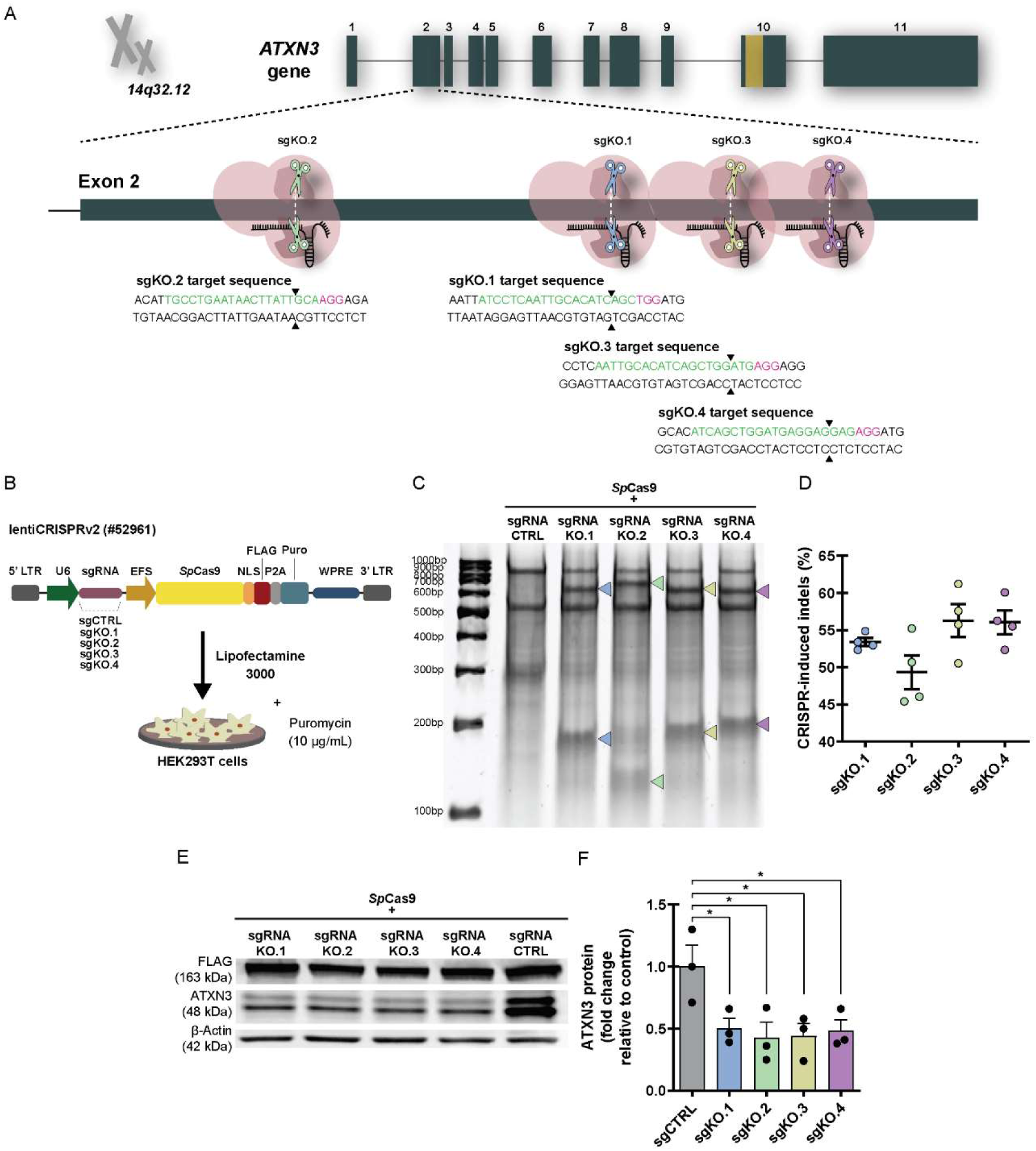
CRISPR-Cas9 system designed to permanently inactivate the human *ATXN3* gene decreases ATXN3 protein expression in HEK293T cells. **(A)** Four guide sequences (sgKO.1, sgKO.2, sgKO.3, and sgKO.4) were designed to recognize exon 2 of the human *ATXN3* gene (sgRNA target sequences are displayed in green). *Sp*Cas9 is recruited to the *locus* of interest, mediating the insertion of a DSB (colored scissors in the scheme and arrowheads in the target sequences) at approximately 3 base pairs upstream of a PAM sequence (sequence displayed in magenta). Subsequently, genome editing is achieved via NHEJ repair pathway for the permanent blocking of *ATXN3* gene expression. **(B)** sgRNA sequences were cloned into a lentiviral expression vector (lentiCRISPRv2, addgene plasmid #52961), which also codifies for a FLAG-tagged *Sp*Cas9 and a puromycin resistance cassette. For the validation of the guide sequences, HEK293T cells were transfected with each of the generated plasmids and maintained in culture for 72 hours (selection medium with puromycin 10 µg/mL for 48 hours). Cells transfected with a guide sequence targeting the bacterial *lacZ* gene (sgCTRL) were used as a negative control. **(C-D)** Locus modification efficiencies were analyzed using Surveyor nuclease assay. Lane 1: DNA ladder (GeneRuler 100 bp, Thermo Fisher Scientific); Lane 2: Cells transfected with the sgCTRL construct; Lanes 3-6: Cells transfected with the sgRNA knock-out guide sequences (sgKO.1, sgKO.2, sgKO.3 and sgKO.4, respectively). Arrowheads indicate the expected DNA fragments, cleaved by Surveyor nuclease. The estimated indel occurrence within human *ATXN3* locus is represented as a percentage (n=4). **(E-F)** Western blot analysis revealed a significant decrease (approximately 0.5-fold change) in ATXN3 protein levels after *Sp*Cas9 targeting of *ATXN3* locus in comparison with the control sequence (n=3). Results are presented as fold change relative to cells transfected with the sgCTRL expressing plasmid. Optical densitometry analysis of ATXN3 fractions were normalized with β-actin and FLAG signals. Statistical significance was evaluated with one-way ANOVA with Dunnett’s post hoc test (*p<0.05). Data are expressed as mean ± SEM. **Abbreviations**: **LTR** (long terminal repeat); **U6** (Pol III promoter); **sgRNA** (single guide RNA); **EFS** (elongation factor 1α short promoter); *Sp*Cas9 (Cas9 nuclease from *Streptococcus pyogenes)*; **NLS** (nuclear localization signal); **FLAG** (FLAG octapeptide tag); **P2A** (2A self-cleaving peptide); **Puro** (puromycin selection marker); **WPRE** (woodchuck hepatitis virus post-transcriptional regulatory element).

Given that all the generated guide sequences successfully modified the *ATXN3* gene, leading to a reduction in ATXN3 protein expression (∼0.5 fold-change), a comprehensive *in silico* analysis was performed to predict potential off-targets for each sgRNA in the human genome. The genomic locations of the predicted off-targets were determined using the UCSC genome browser and mapped to intergenic, intronic and exonic regions (Supplemental Figure S3). Based on this analysis, we selected the sgKO.2 guide for subsequent *in vivo* experiments, as no predicted off-targets (with scores>1, *i.e.* with potential to generate indel mutations) were located in protein-coding exons of the human genome. Co-expression of SpCas9 and sgCTRL or sgKO.2, will be henceforth designated as CRISPR-CTRL and CRISPR-ATXN3 KO, respectively.

### CRISPR-Cas9 system mediates *ATXN3* gene disruption *in vivo* and alleviates neuropathology in a lentiviral-based mouse model of MJD/SCA3

Having demonstrated the efficacy of this gene editing system *in vitro*, we sought to evaluate a viral delivery strategy of CRISPR-Cas9 components into the mammalian brain. Considering the advantages of AAVs over lentivirus, we cloned the previously described sgCTRL and sgKO.2 guide sequences into AAV vectors for *in vivo* delivery. Due to limits related with the packaging capacity of AAVs (∼4.5 kb) ^52^, the *Sp*Cas9 and the sgRNA expressing cassettes were packaged in two distinct viral vectors ^58^.

The delivery efficacy of these constructs was initially tested upon stereotaxic injection of equal amounts of rAAV1/2 particles encoding *Sp*Cas9 (rAAV1/2-*Sp*Cas9) and the *lacZ*-targeting sgRNA (rAAV1/2-sgCTRL) into the striatum of adult mice (Supplemental Figure S4A). Microscopy analysis showed the efficient transduction of rAAV vectors in the brain parenchyma 4 weeks after AAV administration, as can be seen by the detection of the hemagglutinin (HA) tag fused to *Sp*Cas9 (Supplemental Figure S4B) and of the EGFP-KASH fusion protein co-expressed in the sgRNA construct (Supplemental Figure S4C). Importantly, no alterations of DARPP-32 staining were observed, suggesting that neither the rAAV delivery system nor the expression of CRISPR*-*Cas9 components induced toxicity in the adult brain parenchyma (Supplemental Figure S4D).

To evaluate the therapeutic potential of the developed CRISPR-Cas9 system in the context of MJD/SCA3, we stereotaxically co-injected LV-PGK-*ATXN3* 72Q and rAAV1/2 vectors encoding *Sp*Cas9 along with either the sgCTRL (CRISPR-CTRL system, left hemisphere) or the sgKO.2 (CRISPR-*ATXN3* KO system, right hemisphere) into the striatum of adult mice (Figure 4A-B). Immunoblotting of total protein extracts showed an effective reduction of mutant ATXN3 species in CRISPR-*ATXN3* KO transduced striatal tissues, 4 weeks after viral injection (Figure 4C).

**Figure 4.**
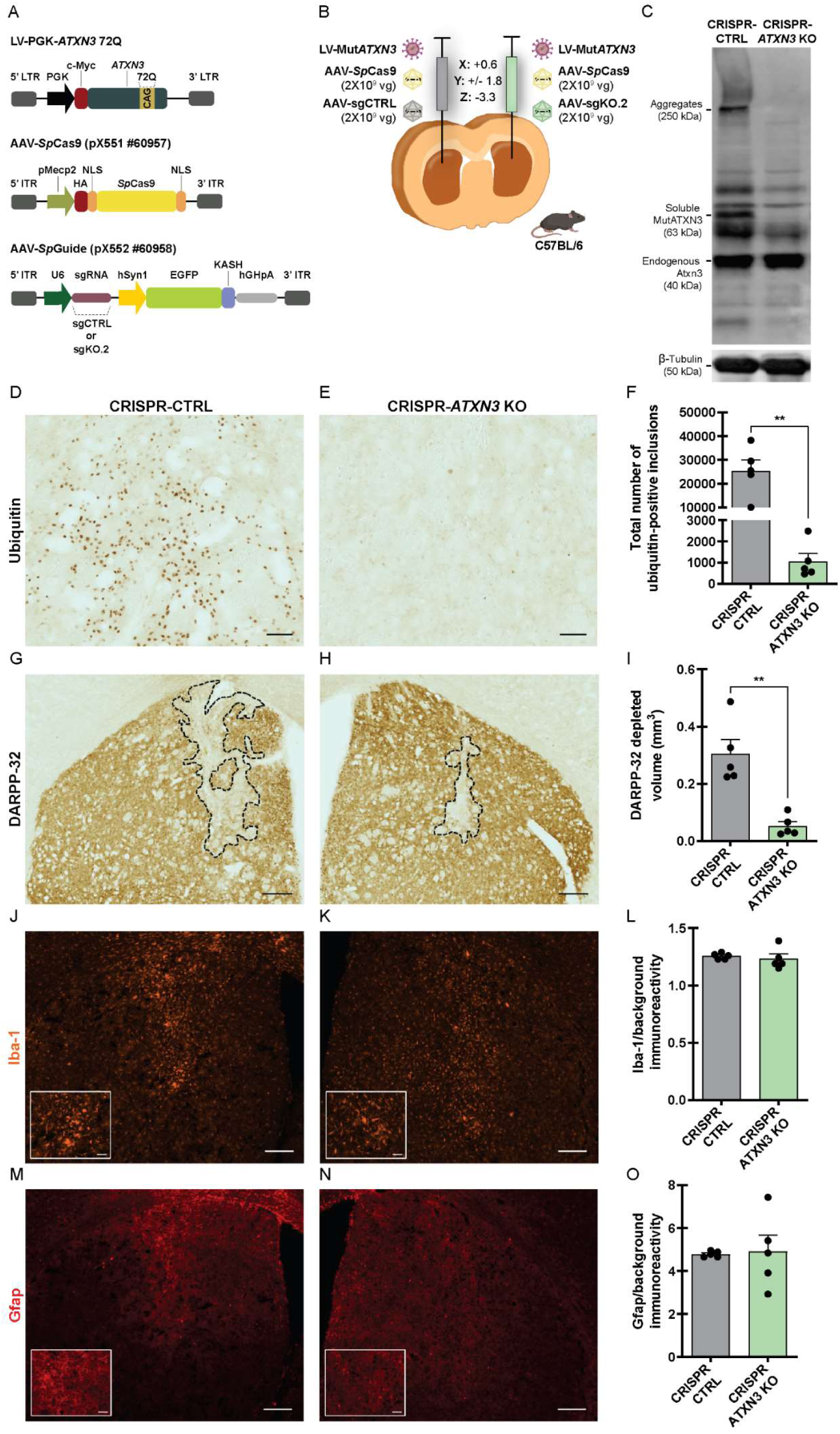
*In vivo* delivery of CRISPR-*ATXN3* KO system alleviates neuropathology in a lentiviral-based mouse model of MJD/SCA3. **(A-B)** Schematic representation of the stereotaxic co-injection of viral vectors in the striatum of C57BL/6 mice. Lentivirus encoding for the human mutant ATXN3 protein with 72Q (Myc-tagged), rAAV1/2 encoding for *Sp*Cas9 (HA-tagged) and rAAV1/2 encoding for the CTRL guide sequence (EGFP-KASH co-expression) were injected in the left hemisphere, serving as experimental control. In the contralateral hemisphere rAAV1/2 encoding for the sgKO.2 sequence were injected, along with LV-PGK-*ATXN3* 72Q and the rAAV1/2-*Sp*Cas9. Four weeks after surgery animals were sacrificed. **(C)** Western blot analysis of striatal homogenates demonstrates that CRISPR-*ATXN3* KO system promotes a reduction of mutant ATXN3 species in treated hemispheres, when compared with the contralateral control hemispheres (data not quantified). **(D-E)** Immunohistochemical peroxidase staining upon labelling of striatal sections with anti-ubiquitin antibody, 4 weeks after stereotaxic surgery. Scale bar, 50 µm. **(F)** CRISPR-*ATXN3* KO injected hemispheres display a drastic reduction in the number of ubiquitin-positive inclusions in comparison with the contralateral control hemisphere, injected with CRISPR-CTRL. **(G-H)** Immunohistochemical analysis using anti-DARPP-32 antibody for expanded ATXN3-derived lesion identification. Treated hemispheres, injected with CRISPR-*ATXN3* KO showed a statistically significant reduction of DARPP-32 depleted volume, as quantified in **(I)**. Scale bar, 200 µm. **(J-K)** Iba-1 immunoreactivity in mouse striata. No statistically significant differences are observed between control (J) and CRISPR-edited hemispheres (K), as quantified in **(L)**. Scale bar, 200 µm in general view images and 50 µm in detail magnifications. **(M-N)** Gfap immunoreactivity in mouse striata. No statistically significant differences in the Gfap immunoreactivity are observed between non-edited (M) and CRISPR-edited striata (N), as quantified in **(O)**. Scale bar, 200 µm in general view images and 50 µm in detail magnifications. Statistical significance was evaluated with paired Student’s t-test (**p<0.01, n=5). Data are expressed as mean ± SEM. **Abbreviations**: **ITR** (invert terminal repeat); **pMecp2** (mouse methyl CpG binding protein 2 promoter); **HA** (hemagglutinin tag); **NLS** (nuclear localization signal); ***Sp*Cas9** (Cas9 nuclease from *Streptococcus pyogenes)*; **U6** (Pol III promoter); **sgRNA** (single guide RNA); **hSyn1** (human synapsin 1 promoter); **EGFP** (enhanced green fluorescent protein); **KASH** (Klarsicht, ANC1, Syne Homology nuclear transmembrane domain); **hGHpA** (human growth hormone gene polyadenylation signal).

In agreement, immunohistochemical analysis of coronal sections obtained from injected mice showed a significant reduction in the total number of ubiquitin-positive inclusions in CRISPR-*ATXN3* KO-injected hemispheres compared to the contralateral CRISPR-CTRL hemispheres (Figure 4D-F; CRISPR-CTRL: 25410 ± 4583 aggregates vs CRISPR-*ATXN3* KO: 1061 ± 371.9 aggregates). Importantly, DARPP-32 immunostaining showed a robust decrease in the depletion volume following *ATXN3* gene disruption (Figure 4G-I; CRISPR-CTRL: 0.30 ± 0.05 mm^3^ vs CRISPR-*ATXN3* KO: 0.05 ± 0.02 mm^3^), demonstrating the ability of the CRISPR-*ATXN3* KO system to preserve neuronal integrity. No statistically significant differences were observed between treated and non-treated hemispheres regarding Iba-1 (Figure 4J-L) or Gfap (Figure 4M-O) immunoreactivity.

Overall, these results suggest that CRISPR-mediated *ATXN3* gene disruption led to a marked amelioration of neuropathological signs in a striatal lentiviral-based mouse model of MJD/SCA3.

### Localized delivery of CRISPR-*ATXN3* KO system reduces intranuclear ATXN3 accumulation in deep cerebellar nuclei of transgenic mice

To further validate the therapeutic effect of the engineered CRISPR-Cas9 system, we used the YAC-MJD84 transgenic mice. This model expresses the full-length human *ATXN3* gene with an expanded CAG repeat (84 CAGs), including all its regulatory elements ^59^. Homozygous YAC-MJD84 mice (YAC-MJD84.2/84.2) recapitulate several neuropathological features characteristic of the disease, exhibiting mild motor deficits as early as 6 weeks of age, which progressively worsen over time ^13^. Overall, this model resembles many aspects of MJD/SCA3 in humans and has been widely used in pre-clinical studies to assess the potential of several MJD/SCA3 therapies ^13,20,60,61^.

Sex-matched, adult homozygous YAC-MJD84 mice were injected into the cerebellar parenchyma with either PBS (control group, n=13) or two different doses of rAAV1/2 encoding the CRISPR-*ATXN3* KO system. The low dose group received a total of 8 x 10^9^ vg, while the high dose group received a total of 1.6 x 10^10^ vg (rAAV1/2-sgKO.2 and rAAV1/2-*Sp*Cas9 at a 1:1 ratio, n=12 per dose). Motor performance was assessed 5 weeks prior to vehicle or rAAV1/2 injection (mice with 4 weeks of age, baseline) and then longitudinally 3, 7 and 11 weeks post-injection (mice with 12, 16 and 20 weeks of age, Figure 5A) Mice body weight was monitored weekly from two weeks of age until euthanasia (21 weeks of age; 12 weeks after injection). No statistically significant differences in body weight were observed between the experimental groups (Figure 5B).

**Figure 5.**
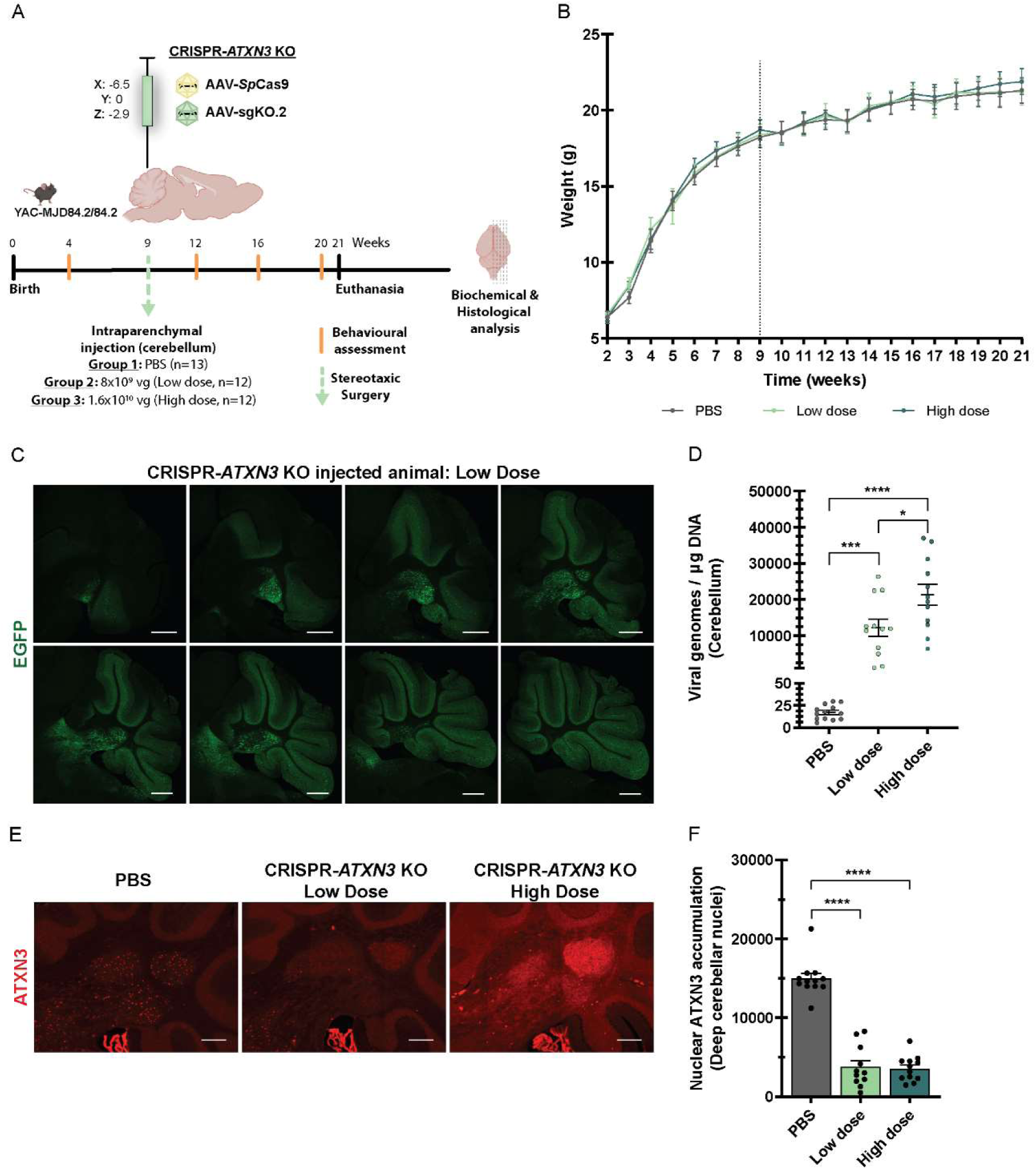
The administration of the CRISPR-*ATXN3* KO system into the cerebellum of adult post-symptomatic transgenic YAC-MJD mice reduces ATXN3 aggregation in the deep cerebellar nuclei. **(A)** Schematic representation of the study design. Three groups of YAC-MJD homozygous mice (YAC-MJD84.2/84.2) underwent stereotaxic surgery at 9 weeks of age for the administration of PBS (n=13), rAAV1/2-CRISPR-*ATXN3* KO low dose (8×10^9^ vg, n=12), or rAAV1/2-CRISPR-*ATXN3* KO high dose (1.6×10^10^ vg, n=12) into the cerebellar parenchyma (coordinates from bregma, X: - 6.5; Y: 0; Z: −2.9). Behavioral assessment was performed throughout the study at 4, 12, 16 and 20 weeks of age. Animals were sacrificed 12 weeks post rAAV1/2 injection (at 21 weeks of age) and brains were collected and processed for biochemical analysis and histological examination. **(B)** Longitudinal body weight assessment (in grams) of homozygous YAC-MJD84.2/84.2 mice used in the study. Each animal was weighed weekly from the second to the twenty-first week of life. No significant differences are observed between the three experimental groups (PBS, low dose, and high dose). Dashed line indicates the week in which the animals underwent stereotaxic surgery. **(C)** Fluorescent immunohistochemical analysis of sagittal brain slices 12 weeks after the injection of the CRISPR-*ATXN3* KO system (low dose) centrally in the cerebellum. The injected rAAV1/2 vectors were efficient to drive the expression of the EGFP-KASH fusion protein encoded in the sgKO.2 construct throughout the cerebellum, including into the deep cerebellar nuclei (DCN). Scale bar, 500 µm. **(D)** Quantification of viral genomes in the cerebella of injected animals (vg/µg of DNA). Statistically significant differences are observed between the three groups of animals. **(E)** Fluorescent immunohistochemical analysis, using an anti-ATXN3 (1H9) antibody on sagittal brain slices of homozygous animals, 12 weeks after the cerebellar injection with PBS (left panels), rAAV1/2-CRISPR-*ATXN3* KO low dose (middle panels) and rAAV1/2-CRISPR-*ATXN3* KO high dose (right panels). Scale bar, 200 µm. **(F)** A statistically significant reduction in the number of cells with intranuclear ATXN3 accumulation is observed in the DCN of homozygous mice injected with both doses of the CRISPR-*ATXN3* KO system, when compared with the PBS-injected animals. Mixed effects analysis with post-hoc Tukey’s multiple comparison test was performed in (B), n=13 for the control group and n=12 for the treatment groups. One-way ANOVA analysis followed by Tukey’s multiple comparisons test was performed in (D), n=13 for the control group and n=12 for the treatment groups. One-way ANOVA analysis followed by Tukey’s multiple comparisons test was performed in (F), n=12 for the control group, n=11 for the low dose group and n=12 for the high dose group. Statistical significance is indicated as *p<0.05, ***p<0.001, ****p<0.0001. Data are expressed as mean ± SEM.

To evaluate rAAV1/2 transduction, sagittal brain slices were stained with an anti-GFP antibody that recognizes the EGFP-KASH fusion protein encoded in the sgKO.2 construct. Although the injection was performed centrally into the cerebellar parenchyma, a widespread and robust diffusion of viral particles was observed throughout the cerebellum, including the deep cerebellar nuclei (DCN), 12 weeks after viral delivery (Figure 5C). Viral genome copies were also quantified in the cerebellum of injected animals. As expected, significant differences were observed in the number of viral copies between the three groups (Figure 5D; PBS-injected group: 17.11 ± 2.261 vg/µg; Low dose group: 12190 ± 2361 vg/µg; High dose group: 21326 ± 2894 vg/µg).

As a consequence of gene editing, a significant reduction in the number of cells with intranuclear ATXN3 accumulation was observed in the DCN of treated animals, a brain region particularly vulnerable in MJD/SCA3 ^10^ (Figure 5E-F; PBS-injected group: 14943 ± 665.3; Low dose group: 3780 ± 788.4; High dose group: 3553 ± 463.9 number of cells with intranuclear ATXN3 accumulation). Importantly, both the low and high dose groups exhibited an approximately 75% reduction in mutant ATXN3 accumulation in the DCN compared to vehicle-injected animals (Figure 5F).

To evaluate the therapeutic effect of the developed strategy, locomotor and exploratory activities were measured using the open-field test (Supplemental Figure S5). Despite a 75% reduction in ATXN3 accumulation in the DCN, no significant improvements were observed in the rAAV-receiving groups across any of the assessed parameters (mean velocity, total distance travelled, time moving fast, resting time, Supplemental Figure S5A-D) at any of the evaluated time points.

### Pre-symptomatic administration of CRISPR-*ATXN3* KO system in the CNS of transgenic mice prevents MJD/SCA3 progression

Following the previous experiment, we hypothesized that the lack of improvement in motor performance could be either due to the inefficient delivery of the therapy to other brain regions affected in the disease and/or due to the late therapeutic intervention (*i.e.* 9 weeks of age), as post-symptomatic treatments may be less effective in reversing motor deficits.

To investigate whether an earlier and broader delivery of the developed system throughout the CNS could ameliorate the motor performance, we administered rAAVs into the cerebrospinal fluid (CSF) of neonatal YAC-MJD84.2/84.2 mice via intracisterna magna (ICM) injections. Transgenic mice received injections of either PBS (vehicle-injected animals) or a total of 1.5×10^10^ viral genomes of rAAV1/2-CRISPR-*ATXN3* KO (rAAV1/2-sgKO.2 and rAAV1/2-*Sp*Cas9 at a 1:1 ratio) between the first and third days of life (P1-P3). The general health and body weight of these animals were monitored weekly until euthanasia (17 weeks of age, Figure 6A). The administration of rAAV1/2-CRISPR-*ATXN3* KO caused no visible impact in mice general health and no statistically significant differences in mice body weight were observed between the experimental groups (Figure 6B).

**Figure 6.**
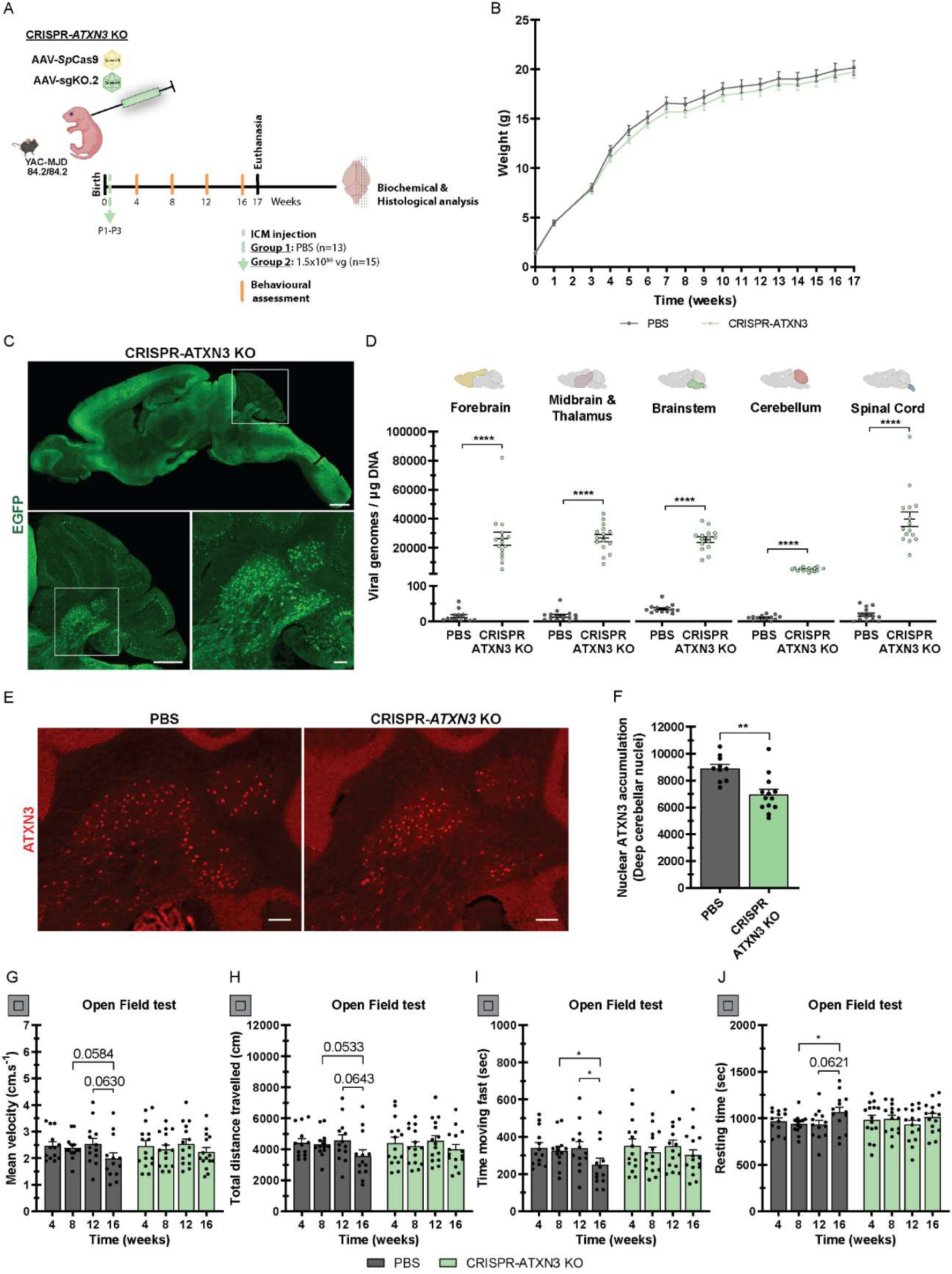
Pre-symptomatic delivery of CRISPR-*ATXN3* KO system directly into the CSF of neonatal homozygous YAC-MJD mice prevents disease progression. **(A)** Schematic representation of the study design. Neonatal homozygous YAC-MJD84 mice received intra-cisterna magna (ICM) administrations of either PBS (n=13) or rAAV1/2-CRISPR-*ATXN3* KO (1.5×10¹⁰ vg, n=15) between the first and third days of life (P1-P3). Behavioral assessment was performed throughout the study at 4, 8, 12 and 16 weeks of age. Animals were sacrificed at 17 weeks of age and brains were collected and processed for biochemical analysis and histological examination. **(B)** Body weight measurements (in grams) were performed weekly from birth until 17 weeks of age. No significant differences are observed between the two experimental groups. **(C)** Fluorescent immunohistochemical analysis of sagittal brain slices shows the widespread expression of EGFP in the brain of transgenic mice, 17 weeks after ICM injections of rAAV1/2-CRISPR-*ATXN3* KO. Scale bar, 1000 µm in the whole brain image (upper panel), 500 µm in the cerebellum image (lower left panel) and 100 µm in the DCN magnification (lower right panel). **(D)** Vector copy distribution throughout the CNS, 17 weeks after ICM viral delivery. Quantifications were performed in five brain regions (forebrain, midbrain and thalamus, brainstem, cerebellum and spinal cord) and are expressed as vg/µg of tissue DNA. **(E)** Fluorescent immunohistochemical analysis, using an anti-ATXN3 (1H9) antibody on sagittal brain slices of YAC-MJD84.2/84.2 animals, 17 weeks after the ICM injection with either PBS (left panel) or rAAV1/2-CRISPR-*ATXN3* KO (right panel). Scale bar, 100 µm. **(F)** A statistically significant reduction in the number of cells with intranuclear ATXN3 accumulation is observed in the DCN of homozygous mice injected with the CRISPR-*ATXN3* KO system, when compared with the PBS-injected animals. **(G-J)** Locomotor horizontal activity over a 30-minute trial in open field test. A disease progression is observed over time in untreated animals given by their reduced locomotor activity. A tendency for a decreased (G) mean velocity (in cm.s⁻¹) and (H) total distance travelled (in centimeters) is observed between 8 and 16 weeks of age and between 12 and 16 weeks of age. A statistically significant reduction of the (I) time (in seconds) moving fast (above 5 cm.s⁻¹), is also observed in these animals, between 8 and 16 weeks of age and between 12 and 16 weeks of age. Moreover, a statistically significant increase in the (J) resting time (in seconds) is observed between 8 and 16 weeks of age, and a tendency for an increased resting time between 12 and 16 weeks of age. No differences are observed in CRISPR-*ATXN3* KO treated mice across the behavioral assessment. Mixed effects analysis with post-hoc Sidak’s multiple comparison test was performed in (B), n=13 for the control group (PBS) and n=15 for the treatment group (CRISPR-*ATXN3* KO). Unpaired Student’s t-test was performed in (D), n=13 for the control group and n=15 for the treatment group, and in (F), n=10 for the control group and n=13 for the treatment group. Two-way ANOVA followed by Tukey’s multiple comparisons test was performed in (G-J), n=13 for the control group and n=15 for the treatment group. Statistical significance of *p<0.05 is indicated. Data are expressed as mean ± SEM.

The CNS biodistribution of the injected rAAV1/2 was determined 17 weeks after viral delivery by immunohistochemistry (EGFP expression, Figure 6C) and by the quantification of vector DNA genome copies in five brain regions: forebrain, midbrain and thalamus, brainstem, cerebellum and spinal cord (Figure 6D). A widespread expression of EGFP was observed in sagittal brain slices of transgenic mice, confirming that ICM administrations of the rAAV1/2-CRISPR system resulted in a broad transduction of the mouse brain (Figure 6C). Additionally, the quantification of viral genomes showed a strong transduction in all five brain regions analyzed, with the highest levels detected in the spinal cord (mean of 4×10^4^ vg/µg tissue DNA) and the lowest in the cerebellum (mean of 5×10^3^ vg/µg tissue DNA, Figure 6D). Considering that the cerebellum has been established as a vulnerable brain region in both patients and animal models of MJD/SCA3 ^10^, the number of cells with intranuclear ATXN3 accumulation was assessed in the DCN of transgenic mice. A statistically significant reduction in the number of cells with intranuclear ATXN3 accumulation was observed in the DCN of treated animals (Figure 6E-F; PBS: 8907 ± 299.5; CRISPR-ATXN3 KO: 6962 ± 383.5 number of cells with intranuclear ATXN3 accumulation), confirming the efficacy of the developed CRISPR-*ATXN3* KO system in this brain region.

Locomotor activity was assessed by open field test every 4 weeks (at 4, 8, 12 and 16 weeks of age). As expected for this animal model, vehicle-injected animals displayed a progressive worsening of the behavior phenotype, as shown by the analysis of locomotor parameters (Figure 6G-J). Indeed, we observed a decrease in mean velocity (Figure 6G), total distance travelled (Figure 6H), time moving fast (Figure 6I) and an increase in resting time (Figure 6J) in control group at 16 weeks compared to 8 and 12 weeks of age. In contrast, no differences in locomotor activity parameters were observed over time in CRISPR-*ATXN3* KO treated mice, demonstrating the ability of a CRISPR-Cas9-based approach to prevent disease progression when administered at a pre-symptomatic stage in the CSF.

## DISCUSSION

An effective and permanent therapeutic approach is an unmet need encountered in polyQ neurodegenerative disorders, including MJD/SCA3. Gene editing tools have made it possible to precisely and permanently modify several disease-causing genes, offering tremendous potential for the development of new therapeutic agents for a wide range of incurable disorders. In view of the urgency to develop new therapies that can stop or delay MJD/SCA3 progression, we sought to evaluate the potential of two nuclease-based approaches (TALENs and CRISPR-Cas9 systems) to mediate the silencing of the human *ATXN3* gene, the gene implicated in MJD/SCA3. A side-by-side comparison revealed that, although both TALEN and CRISPR-Cas9 systems effectively promoted *ATXN3* gene editing, CRISPR-Cas9 system demonstrated the added advantage of preserving neuronal integrity in a lentiviral-based mouse model of the disease. Notably, the developed CRISPR-Cas9 system also proved effective in stalling disease progression in a transgenic YAC-MJD84 mouse model, when administered into the CSF in a pre-symptomatic stage.

Several studies have demonstrated that the non-allele selective silencing of *ATXN3* transcripts in the adult brain is associated with improvements in MJD/SCA3 pathology without any deleterious effects ^13,20,62,63^. Additionally, although some *in vitro* studies showed that ATXN3 is indispensable for neuronal differentiation and cytoskeletal organization ^64,65^, mice and *C. elegans* lacking the codifying gene are viable and phenotypically normal ^66,67^, suggesting that ATXN3 may not be required for proper adult brain function. Therefore, since strategies based on the selective inhibition of the mutant allele will not be appropriate for every patient, these studies indicate that non-allele specific strategies may be used as potential therapeutic approaches, at least before the occurrence of severe neuronal damage.

Targeted gene knock-outs can be generated upon the introduction of a DSB in a codifying exon near the 5’-end of the gene, followed by NHEJ-induction of frameshift mutations and the consequent appearance of an early stop codon ^68,69^. To explore the possibility of using NHEJ to permanently inactivate the human *ATXN3* gene we designed TALEN and CRISPR-Cas9-based systems to target exon 2 of this gene, with no discrimination between the mutant and wild-type alleles.

Three TALEN pairs (TALENs A/D, TALENs B/E and TALENs C/F) were initially designed and constructed. Based on the observation that TALEN pair B and E produced the highest reduction in ATXN3 protein levels in a human cell line, this pair of TALENs was selected for further experiments in a lentiviral-based mouse model. *ATXN3* gene disruption was successfully achieved in the adult brain upon TALENs B/E administration, leading to the consequent decrease in ubiquitin-positive inclusions in treated striata. Nonetheless, we observed microglial and astrocytic activation, evidenced by the elevation of Iba-I and Gfap markers in TALEN-treated hemispheres. In agreement, an increased loss of DARPP-32 immunoreactivity was also observed in treated striata, suggesting that TALENs B/E induce a toxic effect in the brain parenchyma, exacerbating the neuronal dysfunction driven by the expression of the mutant ATXN3 protein.

A side-by-side comparison between zinc finger nucleases (ZFNs) and TALENs *in vitro* showed that TALENs were as effective as ZFNs, but less cytotoxic and more specific, given the significantly reduced nuclease-associated off-target effects ^70,71^. Additionally, numerous clinical trials using TALENs have been approved for *ex vivo* and *in vivo* applications ^72,73^ and, to our knowledge, none of the reports showed the lack of safety of these platforms. Here, it remains unclear whether the observed toxicity is due to off-target effects, specific cellular responses to DSBs, inflammatory responses to the constitutive expression of bacterial proteins in the mouse brain (DNA binding and cleavage domains originally from *Xanthomonas* and *Flavobacterium okeanokoites*, respectively) ^74^, or other underlying causes, which would warrant further investigations before considering the use of TALENs as a potential therapeutic approach for MJD/SCA3.

With the advent of CRISPR-Cas9 systems, a clear change in the preferred genome editing tool was denoted ^75^. This bacterial system, adapted for genome editing in eukaryotic cells, demands the co-expression of an endonuclease Cas9 and a sgRNA molecule in the target cells ^33,35^. Apart from the reported high efficiency and the ability of multiplexing, the mode of recognition used by this system significantly facilitates the re-targeting of *Sp*Cas9 nuclease to new DNA sequences, by simply changing the 20 bp guide sequence of the guide RNA ^33^. Thus, due to the simplicity of the CRISPR-Cas9 strategy, we further explored this tool by designing and constructing four sgRNAs (sgKO.1, sgKO.2, sgKO.3, sgKO.4) to target the exon 2 of the human *ATXN3* gene. The designed sequences proved to be effective in reducing the endogenous levels of the ATXN3 protein, in a non-allelic specific manner. Contrarily to what we observed with TALENs, the *ATXN3* gene inactivation driven by the sgKO.2 resulted in the clearance of ubiquitin-positive inclusions in the striata of a lentiviral-based mouse model of MJD/SCA3, while preventing the striatal neuronal dysfunction. These results indicate a superior safety profile of the CRISPR-Cas9-based knock-out system, when compared to the developed TALENs, in the context of MJD/SCA3.

The lentiviral-based mouse model used in this study offers great advantages for the screening of new therapeutic molecules due to its robustness to assess neuropathological features ^54,76,77^, but it lacks an observable behavioral phenotype. Therefore, we further evaluated the potential of the generated knock-out system in the YAC-MJD84.2 mouse model ^59^. In a first approach, rAAV1/2 vectors expressing the CRISPR-*ATXN3* KO system were injected into the cerebellar parenchyma of adult mice. While these rAAV1/2 vectors were efficient to transduce a large extent of the cerebellum, achieving a 75% reduction in the ATXN3 accumulation in the DCN, this was not sufficient to improve the motor deficits of treated animals in the open-field test. Although there is no estimation regarding the percentage of ATXN3 protein reduction required to achieve a phenotypic improvement in patients, a reduction of at least 50% in protein levels in homozygous mice was expected to produce milder and delayed motor impairments, as observed in hemizygous animals ^13^, that harbor only 2 copies of the human gene. The lack of phenotypic improvement might suggest that the motor deficits observed in this animal model may result from the dysfunction in other vulnerable brain regions, rather than solely from cerebellar dysfunction. Nonetheless, there is also the possibility that the administration of this therapeutic strategy in a post-symptomatic stage may not be sufficient to prevent or revert neuronal damage and the consequent motor impairment.

Considering this, we further performed ICM injections of the rAAV1/2-CRISPR-*ATXN3* KO system in neonatal YAC-MJD84.2/84.2 mice (P1-P3) to simultaneously transduce several brain regions affected in the disease and initiate the treatment at early stages. This strategy proved to be effective, as treated mice have preserved their locomotor activity in opposition to untreated animals which showed a deterioration of their motor performance over time.

In future studies further investigation will be needed to better understand the tolerability of *ATXN3* gene edition in animals treated with different rAAV doses and serotypes that can potentially cause a more pronounced therapeutic benefit.

In conclusion, this study demonstrates that targeting exon 2 of the *ATXN3* gene using TALEN or CRISPR-Cas9 systems enables non-allele-specific gene silencing both *in vitro* and *in vivo*, offering a strategy potentially applicable to the entire MJD/SCA3 patient population. Importantly, this study represents the first demonstration of the *in vivo* effectiveness of gene editing approaches in the context of MJD/SCA3. Among the strategies tested, the CRISPR-Cas9 system emerged as a superior therapeutic candidate by efficiently rescuing neuronal dysfunction and neuroinflammation driven by the expression of the mutant ATXN3 protein in a lentiviral-based mouse model of the disease. Notably, the administration of the CRISPR-Cas9 system at a pre-symptomatic stage resulted in the stalling of disease progression, highlighting its potential as a promising treatment for this devastating disorder.

## MATERIAL AND METHODS

### TALENs directed to the human *ATXN3* gene

#### Selection of TALEN target sites

To knock-out the human *ATXN3* gene, three TALEN pairs were designed to target a 40 bp target sequence in intron 1 and exon 2 of this gene (**5’-TN_19_N_14-20_N_19_A-3’** region, where left TALEN targets a **5’-TN_19_-3’** sequence and right TALEN targets the antisense strand of **5’-N_19_A-3’,** N = A, G, T or C). To facilitate FokI dimerization, left and right target sites were designed to be spaced by 14 to 20 bases of each other. As natural TALE-binding sites always begin with a thymine ^29,30^ and its absence decreases TALE-binding efficiency ^78^, this constraint had to be considered while screening potential target sites. These designs were based on the annotated reference sequence for the human *ATXN3* gene available at the National Center for Biotechnology Information (NCBI, Bethesda, MD, USA) database (https://www.ncbi.nlm.nih.gov/gene, accession number NG_008198.2).

#### Construction of TALENs directed to the human ATXN3 gene

For the assembly of custom 20 bp targeting TALENs (**5’-T_0_N_1-18_N_19_-3’**), each human *ATXN3* target sequence was initially divided into subsequences. Notably, the first base in the TALEN-binding site (5’-thymine, **T_0_**) was already encoded in the N-terminus of the TALEN backbone vector, while the last base at the 3’-end of the recognition site (**N_19_**) is always specified by a half repeat – a tandem repeat comprising 20 amino acid residues. The remaining 18 bp (**N_1_-N_18_**) are specified by 18 full monomers (34 amino acids each), where the residues at positions 12 and 13, known as RVDs, dictate binding specificity according to the following code: asparagine-isoleucine (NI) = A, histidine-aspartic acid (HD) = C, asparagine-glycine (NG) = T, asparagine-histidine (NH) = G ^27–30^. To facilitate assembly, the N_1_-N_18_ sequences were divided into three hexameric subsequences (N_1_-N_6_, N_7_-N_12_ and N_13_-N_18_, see Supplemental Table 1). These hexamers were circularized using a Golden Gate digestion-ligation strategy, following a previously published protocol ^23^.

Each circularized hexamer was PCR-amplified using the previously described primers Hex-F and Hex-R ^23^. To verify amplification, 1.5 µL of each PCR product were electrophoresed in a 2% agarose gel in 1x Tris-borate-ethylenediaminetetraacetic acid (TBE, Bio-Rad, Hercules, CA, USA) buffer, adjacent to a 1 kb Plus DNA ladder (Invitrogen, Thermo Fisher Scientific, Waltham, MA, USA) and stained with SYBR Safe DNA stain (Invitrogen, Thermo Fisher Scientific). The resulting PCR amplicons, visible as 650 bp bands on the gel (Supplemental Figure S1A), were then purified using the QIAquick PCR purification kit (Qiagen, Hilden, Germany), following the manufacturer’s instructions.

Two variants of the FokI nuclease (ELD:KKR mutants, Q486E, I499L, N496D and E490K, I538K, H537R) were used in the context of this work to minimize potential off-target cleavage ^44,45^. All left TALENs recognizing the sense DNA strand were inserted into the FokI-ELD-codifying plasmid, while right TALENs, recognizing the antisense strand, were inserted into the FokI-KKR backbone. Hexamers, half-repeat sequences and the appropriate TALEN backbones were assembled in a second Golden Gate digestion-ligation, following a previously published protocol ^23^. Ligation products were transformed into One Shot Stbl3 chemically competent cells *E. coli* (Invitrogen, Thermo Fisher Scientific), following the manufacturer’s instructions. Each transformation was spread on a pre-warmed selective plate (100 µg/mL ampicillin, Gibco, Thermo Fisher Scientific) and incubated overnight at 37°C. Four colonies of each of the generated TALENs were inoculated into 4 mL of lysogeny broth medium (LB, Sigma-Aldrich, St. Louis, MO, USA) supplemented with 100 µg/mL ampicillin (Gibco, Thermo Fisher Scientific) and incubated for 8 hours at 37°C with constant shaking (200 rpm). Plasmid DNA was isolated form LB-cultures (2 mL) using the QIAprep Spin miniprep kit (Qiagen), in accordance with the manufacturer’s instructions. Each clone was sequence-verified (Genewiz, Azenta Life Sciences, Plainfield, NJ, USA), using the previously described primers ^23^: TALE-Seq-F1 (sequencing of N_1_-N_6_ monomers), TALE-Seq-F2 (sequencing of N_7_-N_12_ monomers) and TALE-Seq-R1 (sequencing of N_13_-N_19_ monomers).

### Generation of sgRNA constructs for *ATXN3* gene editing

#### Target selection for sgRNAs

The specificity of the CRISPR-Cas9 system is defined by a 20-nucleotide guide sequence within the sgRNA, and therefore the selection of the appropriate target DNA sequence is critical to maximize the efficiency of cleavage. Moreover, considering that the RNA-guided *Sp*Cas9 nuclease targets DNA sites in immediate vicinity of a 5’-NGG PAM sequence (N= A, T, C or G), this constraint had to be considered while screening potential *Sp*Cas9 cleavage sites in the human *ATXN3* gene. To select suitable target sites in exon 2 of the *ATXN3* gene, several bioinformatic tools were used. The top four sgRNA candidates exhibiting the best scores in the largest number of tools were selected: sgKO.1, sgKO.2, sgKO.3, sgKO.4. These designs were based on the annotated reference sequence for the human *ATXN3* gene available at the NCBI database (https://www.ncbi.nlm.nih.gov/gene, accession number NG_008198.2).

The non-targeting control sgRNA sequence used in the framework of this study, targets the *lacZ* gene from *Escherichia coli* (sgCTRL: TGCGAATACGCCCACGCGAT) and has previously been described elsewhere ^58^.

#### Cloning of sgRNA sequences in a lentiviral backbone for the co-expression of sgRNA and SpCas9

The lentiCRISPRv2 expression plasmid ^79^ (plasmid #52961, Addgene, Watertown, MA, USA) includes a chimeric sgRNA-codifying region with a cloning site for the specific 20-nucleotide guide sequences immediately upstream of an invariant scaffold. Sequences sgCTRL, sgKO.1, sgKO.2, sgKO.3 or sgKO.4 were cloned in this backbone for *in vitro* experiments. Briefly, this vector was initially digested and dephosphorylated with BsmBI (Esp3I, Thermo Fisher Scientific) and the resulting product was purified upon electrophoretic separation, using the NucleoSpin Gel and PCR clean-up kit (Macherey-Nagel, Düren, Germany). Pairs of partially complementary oligonucleotides (top and bottom, Supplemental Table 2) encoding the guide sequences and containing overhangs matching the BsmBI digested plasmid (Supplemental Table 2, green letters) were synthetized (Invitrogen, Thermo Fisher Scientific). Importantly, since the U6 RNA polymerase III promoter used to express the sgRNA requires a guanine nucleotide at the 5’ terminus of the codifying strand, an extra guanine was included in all cases (Supplemental Table 2, blue letters). Each pair of oligonucleotides was phosphorylated and annealed, following a previously published protocol ^79^. The annealed oligonucleotide pairs were finally ligated to the open vector using the T7 DNA ligase (New England Biolabs, Ipswich, MA, USA) at room temperature for 10 minutes. The ligation product was transformed into One Shot Stbl3 chemically competent *E. coli* (Invitrogen, Thermo Fisher Scientific), following the manufacturer’s instructions. Surviving colonies were selected and inoculated into LB medium (Fisher BioReagents, Pittsburgh, PA, USA), supplemented with 100 µg/mL ampicillin (Enzo Life Sciences, Farmingdale, NY, USA) for 8 hours at 37°C with constant shaking (200 rpm). Plasmid DNA was isolated from each culture using a NZYMiniprep kit (NZYTech, Lisbon, Portugal), following the manufacturer’s instructions.

The correct insertion of each sgRNA-codifying oligonucleotide pair was confirmed by Sanger sequencing (GATC Biotech, Konstanz, Germany), using a primer designed to the U6 promoter (primer LKO.1 5’: 5’-GACTATCATATGCTTACCGT-3’, sequence available at Addgene). The evaluation of the integrity of the long-terminal repeats (LTRs) was performed by restriction analysis using HindIII (Thermo Fisher Scientific), followed by electrophoresis in a 1% agarose gel prepared with 1x Tris-acetate-ethylenediaminetetraacetic acid buffer (TAE, Sigma-Aldrich) and resolved at 80 Volts for 45 minutes.

#### Preparation of sgRNAs for adeno-associated viral delivery

Due to constraints related with AAV packaging capacity, a two-vector system was adopted for *in vivo* applications ^58^: i) AAV-*Sp*Cas9 (vector pX551, plasmid #60957, Addgene) and ii) AAV-*Sp*Guide (vector pX552, plasmid #60958, Addgene). The AAV-*Sp*Guide backbone includes a SapI (LguI, Thermo Fisher Scientific) restriction site upstream of the sgRNA scaffold and was used to clone the sequences sgCTRL and sgKO.2 for the knock-out experiments in the adult brain. The overhangs for the ligation were properly changed in each oligonucleotide to match the SapI digested plasmid (Supplemental Table 2, in green), while maintaining the 20-nucleotide target sequences and the appended 5’-guanine required by the U6 promoter (Invitrogen, Thermo Fisher Scientific, Supplemental Table 2, in blue). After the digestion of the AAV-*Sp*Guide vector with the SapI (LguI, Thermo Fisher Scientific) restriction enzyme, the resulting product was purified upon electrophoretic separation, using the NucleoSpin Gel and PCR clean-up kit (Macherey-Nagel). Each pair of oligonucleotides was phosphorylated, annealed and ligated, following previously published protocols ^58^. The ligation product was transformed into recombination-deficient bacteria (One Shot Stbl3, Invitrogen, Thermo Fisher Scientific). Surviving colonies were then selected and inoculated into LB medium (Fisher BioReagents) supplemented with ampicillin (100 µg/mL, Enzo Life Sciences). Plasmid DNA was isolated from each culture using the NZYMiniprep kit (NZYTech) and sequenced (GATC Biotech) with the U6-forward primer (primer LKO.1 5’: 5’-GACTATCATATGCTTACCGT-3’, available at Addgene) to confirm the correct insertion of the sgCTRL and sgKO.2-codifying oligonucleotide pairs.

The evaluation of the integrity of the invert terminal repeats (ITRs) was performed by restriction analysis using SmaI (Thermo Fisher Scientific), followed by electrophoresis in a 1% agarose gel in 1x TAE buffer (Sigma-Aldrich) and resolved at 80 Volts for 45 minutes.

### Cell culture and transfection: functional validation of TALENs and sgRNA sequences designed to target the human *ATXN3* gene

Human embryonic kidney 293 cells, stably expressing the SV40 large T antigen (HEK293T) were cultured in Dulbecco’s Modified Eagle Medium (DMEM, Sigma-Aldrich) high glucose, supplemented with 10% fetal bovine serum (FBS; Gibco, Thermo Fisher Scientific) and 1% penicillin/streptomycin (Gibco, Thermo Fisher Scientific) at 37°C under a humidified atmosphere containing 5% CO_2_.

For transfection experiments, 2.75 x 10^5^ cells were plated per well (in 12-well plates) the day before transfection. Lipofectamine 3000 reagent (Invitrogen, Thermo Fisher Scientific) was used for plasmid DNA transfection. Complex formation was performed by combining a mixture of Lipofectamine-Opti-MEM reduced serum media (Gibco, Thermo Fisher Scientific) and DNA-Opti-MEM-P3000 Reagent (Invitrogen, Thermo Fisher Scientific) in a 1:1 ratio, in accordance with the manufacturer’s instructions. Initially, Lipofectamine 3000 was diluted in Opti-MEM and added to a second solution containing the plasmid DNA diluted in Opti-MEM and P3000 reagent. The mixture was incubated at room temperature for 5 minutes and added dropwise to cell cultures. The medium was completely replaced 4 hours later.

To assess TALENs’ functionality we co-transfected HEK293T cells with 1.6 µg of the generated TALEN pairs in technical triplicates: AAV-TALEN A and AAV-TALEN D; AAV-TALEN B and AAV-TALEN E; AAV-TALEN C and AAV-TALEN F. Cells transfected with the transfection reagent alone (mock transfection) were used as controls. Cell collection was performed 48 hours post-transfection, for either genomic DNA purification or protein isolation.

The assessment of sgRNAs’ functionality was performed upon transfection of HEK293T cells with 1.6 µg of the generated plasmids in technical triplicates: LV-*Sp*Cas9-sgKO.1, LV-*Sp*Cas9-sgKO.2, LV-*Sp*Cas9-sgKO.3 or LV-*Sp*Cas9-sgKO.4. Cells transfected with LV-*Sp*Cas9-sgCTRL were used as controls. Twenty-four hours after transfection cell medium was completely substituted by fresh medium containing 10 µg/mL puromycin (Gibco, Thermo Fisher Scientific) and cultures were maintained for 48 hours in antibiotic selection conditions. Cell collection was performed 72 hours post-transfection, for either genomic DNA purification or protein extract preparation.

### Bioinformatic prediction of potential off-targets

Potential off-target loci for the designed sgKO sequences in the human genome were computationally predicted using the Benchling web tool (https://benchling.com/). This tool is based on a previously designed algorithm ^80^ and takes into consideration that *Sp*Cas9 can mediate the cleavage of genomic off-targets in the presence of either 5’-NGG or 5’-NAG PAM sequences. The screen identifies 50 putative off-targets per guide sequence, scored in accordance with the likelihood of off-target binding. Genomic locations of the predicted off-targets in the human genome were determined using the UCSC genome browser (https://genome.ucsc.edu/).

### Viral production, purification and titer assessment

Lentiviral vectors were produced in HEK293T cells with a four-plasmid system, as previously described ^56^. Human immunodeficiency virus type 1 (HIV-1) vectors were pseudotyped with the vesicular stomatitis virus glycoprotein (VSV-G) envelope and encoded for the human mutant ATXN3 with 72 glutamines (LV-PGK-*ATXN3* 72Q), under the control of phosphoglycerate kinase (PGK) promoter ^53^. Lentiviral particles were concentrated by ultracentrifugation of the culture medium containing virus and resuspended in sterile 0.5% bovine serum albumin (BSA, Merck Millipore, MilliporeSigma, Burlington, MA, USA) in phosphate-buffered saline (PBS) 1x pH=7.4 (Fisher BioReagents). The viral particle content of each batch was determined using the Retro-Tek HIV-1 p24 antigen ELISA kit to detect HIV-1 p24 antigen levels (Gentaur, Brussels, Belgium), following the manufacturer’s instructions.

Mosaic recombinant adeno-associated virus co-expressing capsid proteins from AAV1 and AAV2 (rAAV1/2) were produced in HEK293T cells, following a previously described protocol ^81^. Briefly, this method relies on the triple transfection of cells with plasmids encoding for the AAV1 and AAV2 serotypes, with the adenoviral helper genes (pHelper) and the transgene of interest: single stranded green fluorescent protein (rAAV-ssGFP), rAAV-TALEN B, rAAV-TALEN E, rAAV-*Sp*Cas9, rAAV-sgCTRL and rAAV-sgKO.2. Subsequently, the produced rAAVs are harvested from transfected cells and purified by fast protein liquid chromatography (FPLC), using an ÄKTA pure 25 system (Cytiva, Marlborough, MA, USA). rAAV titer was determined by quantitative real-time PCR using the AAVpro Titration kit (for Real Time PCR) Ver.2 (Takara Bio Inc, Shiga, Japan), following the manufacturer’s instructions.

### Animal handling

All experiments involving animals were carried out in compliance with the European Union Community directive (2010/63/EU), for the care and use of laboratory animals. Additionally, animal procedures were approved by the Responsible Organization for the Animal Welfare of the Faculty of Medicine and Center for Neuroscience and Cell Biology of the University of Coimbra (CNC-UC) licensed animal facility. Researchers received suitable training (Federation of European Laboratory Animal Science Associations (FELASA)-certified course) and certification from Portuguese authorities (Direcção Geral de Alimentação e Veterinária) to perform the experiments. Mice were housed in plastic cages (365 x 207 x 140 mm) with a maximum number of five animals per cage (minimum of two) and maintained in a standard 12 hour light/dark cycle with constant temperature (22 ± 2°C) and humidity (55 ± 15%). Food and water were provided *ad libitum.* All efforts were made to minimize animal suffering.

#### Lentiviral-based mouse model

Six-week-old C57BL/6 male mice were obtained from Charles River Laboratories (Barcelona, Spain). One week after acclimation, C57BL/6 mice were anesthetized by intraperitoneal administration of a mixture of xylazine/ketamine (4/80 mg/Kg body weight; Rompun, Bayer, Leverkusen, Germany/Clorketam 1000, Vétoquinol, Lure, France). Mice were stereotaxically injected with viral vectors into the striatum, using the following coordinates calculated from bregma: anteroposterior: +0.6 mm; lateral: ±1.8 mm; ventral: −3.3 mm. Injections were performed using a 30 G blunt-tip injection needle (7803-07, Hamilton Company, Reno, NV, USA) connected to a 10 µL Hamilton syringe (7653-01, Hamilton Company) at an infusion rate of 0.5 µL/min (injection of rAAV vectors alone), or 0.25 µL/min (injection of mixtures containing rAAVs and lentiviral particles). After the infusion, the syringe was allowed to remain in place for 5 minutes and then retracted 0.3 mm, where it was held for an additional 2 minutes prior to its complete removal, to allow viral diffusion while minimizing backflow. Concentrated viral stocks were thawed on ice and diluted using sterile PBS 1x pH=7.4 (rAAV vectors alone) or 0.5% BSA/PBS (for mixtures containing lentiviral vectors).

For the assessment of rAAV1/2 transduction efficiency in the brain parenchyma, mice received a single 2 µL injection of i) a mixture of rAAV1/2-TALEN B and rAAV1/2-TALEN E (1:1) in the left hemisphere containing a total of 4×10^9^ viral genomes (vg) or ii) rAAV1/2-*Sp*Cas9 and rAAV1/2-sgCTRL, premixed in equal amounts, in the left hemisphere (4×10^9^ vg total). For the evaluation of TALENs’ editing efficiency, animals were bilaterally co-injected with concentrated lentiviral vectors, LV-PGK-*ATXN3* 72Q (300ng of p24 antigen) and i) rAAV1/2-ssGFP (4×10^9^ vg) in the left control hemisphere or ii) a mixture of 1:1 ratio of rAAV1/2-TALEN B (2×10^9^ vg) and rAAV1/2-TALEN E (2×10^9^ vg) in the right hemisphere, in a total volume of injection of 2 µL. For the evaluation of the editing efficiency of the generated CRISPR-*ATXN3* KO system, animals were bilaterally co-injected with i) concentrated lentiviral vectors, LV-PGK-*ATXN3* 72Q (300 ng of p24 antigen), ii) rAAV1/2-*Sp*Cas9 (2×10^9^ vg) and iii) rAAV1/2-sgCTRL (2×10^9^ vg, left hemisphere) or rAAV1/2-sgKO.2 (2×10^9^ vg, right hemisphere), in a total volume of 2 µL.

After surgery mice were maintained in their home cages and received post-operative care for at least 7 days. Animals were sacrificed 4 weeks later for tissue processing.

#### YAC-MJD84 transgenic mice

Hemizygous YAC-MJD84 (YAC-MJD84.2) transgenic mice expressing the full-length human ATXN3 gene with 84 CAG repetitions ^59^ were acquired from the Jackson Laboratory (Bar Harbor, ME, USA). A colony of these transgenic mice was established and maintained in the animal facility at the CNC-UC.

Genotyping was performed using genomic DNA isolated from an ear punch prior to weaning (at 10 days of age) and confirmed postmortem. Genomic DNA was extracted from ears using the GeneJET Genomic DNA purification kit (Thermo Fisher Scientific), accordingly with the manufacturer’s instructions. YAC-MJD84 transgenic mice hemizygosity/homozygosity was determined by quantitative real-time PCR using TaqMan Fast Advanced Master Mix (Applied Biosystems, Thermo Fisher Scientific) to amplify a fragment of the human *ATXN3* transgene (forward primer: CCTCAATTGCACATCAGCTGGAT; reverse primer: AACGTGCGATAATCTTCACTAGTAACTC; probe sequence: CTGCCATTCTCATCCTC, Thermo Fisher Scientific), which was normalized for the mouse *Beta Actin* (Cat # Mm02619580_g1, Thermo Fisher Scientific). Primer and probe sequences were described elsewhere ^82^. The reactions were performed in a CFX96 Real-Time PCR detection system (Bio-Rad). In this study, only homozygous YAC-MJD84 (YAC-MJD84.2/84.2) transgenic mice were used. The number of CAG repeats in the human *ATXN3* gene from YAC-MJD84.2/84.2 mice was determined by fragment length analysis, following a previously optimized protocol ^83^. Only homozygous animals expressing a minimum of (CAG)_73_ repetitions were included in these studies.

##### YAC-MJD84 transgenic mice: Stereotaxic cerebellar viral delivery

Stereotaxic administration of CRISPR-*ATXN3* KO system or vehicle into the cerebellum was performed on nine week-old YAC-MJD84.2/84.2 mice under vaporized isoflurane anesthesia, as previously described ^81^. Twelve to thirteen sex-matched YAC-MJD84.2/84.2 mice were included per experimental group: i) PBS (vehicle control group, n=13), ii) rAAV1/2-CRISPR-*ATXN3* KO low dose (8×10^9^ vg, rAAV1/2-sgKO.2 and rAAV1/2-*Sp*Cas9 at a 1:1 ratio, n=12), or iii) rAAV1/2-CRISPR-*ATXN3* KO high dose (1.6×10^10^ vg, rAAV1/2-sgKO.2 and rAAV1/2-*Sp*Cas9 at a 1:1 ratio, n=12). Concentrated viral stocks were thawed on ice and diluted using sterile PBS 1x (pH=7.4). Mice received a single injection of 4.25 µL centrally into the cerebellar parenchyma, using the following coordinates calculated from bregma: anteroposterior: −6.5 mm; lateral: 0 mm; ventral: −2.9 mm. Injections were performed using a 30 G blunt-tip injection needle (7803-07, Hamilton Company) connected to a 10 µL Hamilton syringe (7653-01, Hamilton Company) at an infusion rate of 0.5 µL/min. After the infusion, the syringe was allowed to remain in place for 3 minutes and then slowly retracted 0.3 mm, where it was held for an additional 2 minutes prior to its complete removal from the mouse brain, in order to allow viral diffusion and minimize backflow. After surgery, mice were maintained in their home cages and received post-operative care for at least 7 days. Animals were sacrificed 12 weeks later for tissue processing.

##### YAC-MJD84 transgenic mice: Neonatal intra-cisterna magna injections of viral vectors

To evaluate the potential therapeutic benefit of the generated CRISPR-*ATXN3* KO system in a pre-symptomatic stage, neonatal homozygous YAC-MJD84 mice received an intra-cisterna magna (ICM) injection of either PBS or rAAV1/2-CRISPR-*ATXN3* KO. A total of 15 pups were injected in both control (n=7 females, n=8 males) and treated groups (n=10 females, n=5 males). During the course of the study, two animals from the control group (1 male and 1 female) were euthanized due to overgrowth of incisors. As a result, this experimental group was reduced to 13 animals. The administration of rAAV1/2-CRISPR-*ATXN3* KO caused no apparent impact in mice general health.

Briefly, pregnant females were housed singly and monitored daily from embryonic day 17 to 21, with the least possible disturbance, to ensure that new-born pups could be dosed on postnatal days 1 to 3 (P1-P3). After weighting, pups were placed on ice for cryo-anesthetization, for approximately 1 min. Viral preps containing a total of 1.5×10^10^ vg of rAAV1/2-sgKO.2 and rAAV1/2-*Sp*Cas9 (1:1 ratio, total volume of 3.19 µL), were manually injected into the cisterna magna using a 5 µL Neuros syringe, connected to a 33 G beveled tip needle (65460-03, Hamilton Company), with the sleeve adjusted to 1.5 mm depth. The needle was left on place for a few seconds after the injection. Each pup was identified by tattooing of the paws, carefully cleaned and allowed to recover in a pre-warmed pad. Pups were returned to their original cage, after being rubbed with their original bedding to prevent mother’s rejection. Injected animals were sacrificed 17 weeks later for tissue processing.

### Behavioral assessment: Open-field motor evaluation

Locomotor activity of mice was assessed by placing the animals in the center of a photobeam activity system open-field apparatus (45 cm x 45 cm arena; IR Actimeter system, Panlab SA, Barcelona, Spain) and mice activity was recorded for 30 minutes with the Acti-Track system (Panlab SA). During the course of the experiment the operator was outside the experimental room. Four parameters were evaluated from the recorded data: i) Mean velocity of mice (measured in cm.s^-1^), ii) total distance travelled during the experimental time (measured in centimeters), iii) total time moving fast (measured in seconds), *i.e.* time spent by the mice moving above the threshold of 5 cm.s^-1^ and iv) total resting time (measured in seconds). Motor assessments of YAC-MJD84.2/84.2 mice were performed at 4, 12, 16 and 20 weeks of age (animals injected in a post-symptomatic phase into the cerebellum) or at 4, 8, 12 and 16 weeks of age (animals injected in a pre-symptomatic phase via ICM).

### Brain tissue collection

Mice from the lentiviral-based model were sacrificed 4 weeks after stereotaxic injection with an overdose of xylazine/ketamine (8/160 mg/Kg body weight, intraperitoneal administration). For biochemical analysis, brains were quickly removed from the heads and mice striata were dissected and stored at −80°C until use. For histological analysis, mice were transcardially perfused with ice-cold PBS 1x (pH=7.4), followed by 4% paraformaldehyde (PFA, Sigma-Aldrich). Brains were collected and post-fixed in 4% PFA for 24 hours, at 4°C. Cryoprotection was mediated by the immersion of brains in a 25% sucrose/PBS solution for 48 hours at 4°C. Brains were frozen at −80°C and sliced into coronal sections with 25 µm in thickness, using a cryostat (LEICA CM3050S, Leica Microsystems, Wetzlar, Germany) at −21°C. Slices were collected in anatomical series, being stored at 4°C as free-floating sections in PBS 1x (pH=7.4) supplemented with 0.05% (m/v) sodium azide (Sigma-Aldrich).

Twelve or seventeen weeks after rAAV delivery (at 21 or 17 weeks of age), YAC-MJD84.2/84.2 mice were terminally anaesthetized by intraperitoneal administration of a lethal dose of xylazine/ketamine (8/160 mg/Kg body weight) and transcardially perfused with ice-cold PBS 1x (pH=7.4). Brains were harvested and cut along the middle line in two halves. Left hemispheres were macrodissected in five brain regions and immediately stored at −80°C for DNA extraction: i) forebrain, ii) midbrain and thalamus, iii) brainstem, iv) cerebellum and v) spinal cord. Right hemispheres were post-fixed in 4% PFA for 24 h at 4°C, and embedded in a 25% sucrose/PBS solution for 48 hours at 4°C, for histological assessments. Brains were frozen at −80°C and sliced into sagittal sections with 30 µm in thickness, using a cryostat (CryoStar NX50, Thermo Fisher Scientific) at −21°C. Brain sections were collected in anatomical series as free-floating sections in PBS 1x (pH=7.4) supplemented with 0.05% (m/v) sodium azide and stored at 4°C until further processing.

### Genomic DNA extraction

HEK293T cell cultures were rinsed with PBS 1x (pH=7.4) before genomic DNA extraction. Genomic DNA was extracted using the GeneJET Genomic DNA purification kit (Thermo Fisher Scientific) accordingly with the manufacturer’s instructions. The AllPrep DNA/RNA/Protein mini kit (Qiagen) was used for the extraction of genomic DNA from the different brain regions of YAC-MJD84.2/84.2 mice, following the manufacturer’s instructions. In all cases, genomic DNA concentration and purity were determined using a Nanodrop 2000 Spectrophotometer (Thermo Fisher Scientific).

### Assessment of genome editing: Surveyor nuclease assay

To evaluate the efficiency of TALENs and CRISPR-Cas9 in indel (insertions/deletions) generation a Surveyor nuclease assay (Integrated DNA Technologies, Coralville, IA, USA) was performed ^47^.

The first step of this assay involves genomic PCR amplification of the region surrounding TALEN and CRISPR target sites. Each PCR reaction (50 µl) was prepared on ice, using the following components: 10µl of 5x Phusion HF Buffer (Thermo Fisher Scientific), 1 µl of 10mM deoxynucleotide triphosphates (dNTPs, Thermo Fisher Scientific), forward and reverse primers with a final concentration of 0.5 µM each, 50 ng of DNA template and 0.5 µl (one unit) of Phusion High-Fidelity DNA Polymerase (Thermo Fisher Scientific). Exon 2 of the full-length human *ATXN3* gene was amplified with primer sequences 5’-TTTAGGGGAAGATACTATACAATTC-3’ and 5’-TTTGACAGGTAGTTGAAGCAAG-3’, while the exon 2 of the *ATXN3* cDNA was amplified using primer sequences 5’-ACTAGTGGATCCGTCGACAT-3’ and 5’-CCCGTCAAGAGAGAATTCAAGT-3’. The PCR-amplification was performed in a Veriti 96-Well thermal cycler (Applied Biosystems, Thermo Fisher Scientific) with the following protocol: one cycle at 98°C for 30 seconds (initial denaturation), and 35 cycles at 98°C for 10 seconds (denaturation), X°C for 10 seconds (annealing temperature of 56°C for the genomic *ATXN3* and 62°C for the *ATXN3* cDNA), 72°C for 30 seconds (extension), with a final extension at 72°C for 10 minutes. To check for the specificity of the amplification (single-band products), 2 µL of each PCR product were electrophoresed in 1.5% agarose gels, adjacent to the GeneRuler 100 bp DNA ladder (Thermo Fisher Scientific). Agarose gels were prepared with 1x TAE buffer (Sigma-Aldrich) and resolved at 80 Volts for 45 minutes.

PCR amplicons were purified with the NucleoSpin Gel and PCR clean-up (Macherey-Nagel), according to the manufacturer’s recommendations. DNA heteroduplex formation and the Surveyor nuclease S (Integrated DNA Technologies) digestion were performed in accordance with the recommended parameters ^23,46^. The resulting fragments were separated by electrophoresis on a 10% polyacrylamide (PAGE, Bio-Rad) gel prepared with 1x TBE (Bio-Rad), along with the GeneRuler 100 bp DNA ladder (Thermo Fisher Scientific) or the GeneRuler 50 bp DNA ladder (Thermo Fisher Scientific), at 60 Volts until the bromophenol blue dye had migrated to the bottom of the gel. Gels were post-stained with Gelstar dye (Lonza, Basel, Switzerland) diluted in 1x TBE (1:10,000) for 30 minutes at room temperature and visualized with the Gel Doc EZ imaging system (Bio-Rad). Estimation of cleavage intensity was performed through the measurement of the integrated intensity of the bands corresponding to the undigested PCR amplicons and to the cleaved fragments, using the Image Lab software, version 5.1 (Bio-Rad). Indel frequency was calculated based on a formula described elsewhere ^23,46^. Expected cleaved fragment sizes for *ATXN3*-edited regions are specified in Supplemental Table 3.

### Determination of the number of viral genome copies in animal tissues

The number of viral genomes in brain samples of YAC-MJD84.2/84.2 mice was detected by quantitative real-time PCR using the AAVpro Titration kit (for Real Time PCR) Ver.2 (Takara Bio Inc), following the manufacturer’s instructions. Briefly, DNA samples were initially diluted (final concentration of 10 ng/µL) and subjected to quantitative real-time PCR amplification alongside with serial dilutions of a calibrated control standard. Final result was expressed in terms of rAAV vg/µg of tissue DNA.

### Protein extraction and western blot analysis

After rinsing cellular cultures with PBS 1x (pH=7.4), proteins were extracted from HEK293T cells with a radio-immunoprecipitation assay (RIPA) buffer: 50 mM Tris-base, pH=8.0 (Fisher BioReagents); 150 mM sodium chloride (NaCl, Acros Organics, Antwerp, Belgium); 5 mM ethylene glycol tetraacetic acid (EGTA, Sigma-Aldrich); 1% Triton X-100 (Fisher BioReagents); 0.5% sodium deoxycholate (Sigma-Aldrich); 0.1% sodium dodecyl sulphate (SDS, Fisher BioReagents), supplemented with protease inhibitor cocktail (Roche Diagnostics, Basel, Switzerland), 0.2 mM phenylmethylsulphonyl fluoride (PMSF, Sigma-Aldrich), 1 mM dithiothreitol (DTT, Sigma-Aldrich), 1 mM sodium orthovanadate (Sigma-Aldrich) and 5 mM sodium fluoride (Sigma-Aldrich). The lysate was further sonicated by 2 series of 4 seconds ultra-sound pulses (1 pulse/sec) and centrifuged at 13,400 rcf for 20 minutes at 4°C. The supernatant was collected, and protein concentration determined using Bradford reagent (Bio-Rad). For the extraction of proteins from mice striata (lentiviral-based model), this brain tissue was initially placed on iced patches and sliced into several pieces, using a clean scalpel. Only half of the striatal tissue was lysed with RIPA buffer and homogenized. After sonication (2 series of 4 pulses), protein concentration was assessed using Bradford reagent (Bio-Rad).

Protein samples were denatured with 6x sample buffer (0.5 M Tris-HCl/0.4% SDS, pH=6.8, 9.3% DTT, 10% SDS, 30% glycerol and 0.012% bromophenol blue) and incubated for 5 minutes at 95°C. Thirty to sixty micrograms of total denatured protein (from cells and mice tissue, respectively) were loaded into a SDS-PAGE gel (4% stacking, 10% resolving, Bio-Rad) for subsequent electrophoretic separation. After protein transfer to polyvinylidene difluoride (PVDF) membranes (Merck Millipore, MilliporeSigma), each membrane was blocked in 5% non-fat milk diluted in 0.1% Tween 20 in Tris buffered saline (TBS-T, Fisher BioReagents) for 1 hour at room temperature. The following primary antibodies (diluted in blocking solution) were used for the overnight incubation at 4°C: mouse monoclonal anti-ATXN3 antibody (1H9; 1:5000; Merck Millipore, MilliporeSigma); mouse monoclonal anti-FLAG antibody (M2; 1:1000; Sigma-Aldrich); mouse monoclonal anti-β-actin antibody (AC74; 1:10,000; Sigma-Aldrich); mouse monoclonal anti-β-tubulin 1 antibody (SAP.4G5; 1:10,000; Sigma-Aldrich). After three 15 min-washes in TBS-T, membranes were incubated with the goat anti-mouse alkaline phosphatase-conjugated secondary antibody (1:10,000; Thermo Fisher Scientific Pierce) for 2 hours at room temperature. Membranes were washed (three washes of 15 min each) and revealed with enhanced chemifluorescence substrate (ECF, Cytiva) and protein bands were detected by chemifluorescence imaging (Chemidoc imaging system, Bio-Rad).

Semi-quantitative analysis was carried out based on the optical density of the scanned membranes using Image Lab analysis software (version 5.1, Bio-Rad). Normalization of the total amount of protein loaded in each lane was performed based on β-actin signals. In cellular experiments using the CRISPR-Cas9 system, FLAG signals were also taken into consideration for the normalization of transfection efficiencies.

### Immunohistochemical analysis: Bright-field immunohistochemistry

Free-floating brain coronal sections were incubated for 30 minutes at 37°C with a 0.1% phenylhydrazine (Merck Millipore, MilliporeSigma) /PBS solution, to block endogenous peroxidases. After permeabilization and blocking of nonspecific staining in PBS/0.1% Triton X-100 (Fisher BioReagents) with 10% normal goat serum (NGS, Gibco, Thermo Fisher Scientific) for 1hour at room temperature, sections were incubated overnight at 4°C with primary antibodies diluted in blocking solution: mouse monoclonal anti-FLAG antibody (M2; 1:1000; Sigma-Aldrich), rabbit polyclonal anti-ubiquitin antibody (1:300, Enzo Life Sciences) or rabbit polyclonal anti-dopamine and cyclic AMP-regulated neuronal phosphoprotein 32 (DARPP-32) antibody (1:1000; Merck Millipore, MilliporeSigma). Sections were then washed and incubated with the biotinylated secondary goat anti-mouse antibody (1:250; Vector Laboratories, Newark, CA, USA) or biotinylated secondary goat anti-rabbit antibody (1:250; Vector Laboratories) diluted in blocking solution, for 2 hours at room temperature. Bound antibodies were visualized using the Vectastain ABC kit (Vector Laboratories), using the 3,3’-diaminobenzidine tetrahydrochloride (DAB, Vector Laboratories) as substrate. Sections were then placed in slides previously coated with a gelatine solution, dehydrated with ethanol solutions (with increasing concentrations) and xylene substitute (Sigma-Aldrich) and finally coverslipped with Eukitt quick-hardening mounting medium (Sigma-Aldrich).

Images were captured with the slide scanner Zeiss Axio Scan.Z1 microscope (Carl Zeiss Microscopy GmbH, Oberkochen, Germany), equipped with a Hitachi HV-F202SCL camera, using the Plan-Apochromat 20x/0.8 objective and ZEN software (Carl Zeiss Microscopy GmbH).

### Immunohistochemical analysis: Fluorescence immunohistochemistry

Free-floating brain coronal (lentiviral-based model) or sagittal (YAC-MJD84 model) sections were incubated for 1 hour at room temperature in blocking/permeabilization solution (PBS/0.1% Triton X-100 containing 10% NGS) and then incubated overnight at 4°C with primary antibodies diluted in blocking solution: rabbit polyclonal anti-ionized calcium binding adaptor molecule 1 (Iba-1) antibody (1:1000; Fujifilm Wako Chemicals, Richmond, VA, USA), mouse monoclonal anti-glial fibrillary acidic protein (Gfap) antibody (GA5; 1:500; Merck Millipore, MilliporeSigma), rabbit polyclonal anti-GFP antibody (1:1000; Invitrogen, Thermo Fisher Scientific), mouse monoclonal anti-HA antibody (16B12; 1:1000; Biolegend, San Diego, CA, USA), chicken polyclonal anti-GFP antibody (1:1000, Abcam, Cambridge, United Kingdom) and mouse monoclonal anti-ATXN3 antibody (1H9; 1:5000; Merck Millipore, MilliporeSigma). After a three-washing step, sections were incubated at room temperature for 2 hours, with secondary antibodies coupled to fluorophores, diluted in blocking solution: goat anti-rabbit Alexa Fluor 568, goat anti-mouse Alexa Fluor 647, goat anti-rabbit Alexa Fluor 488, goat anti-mouse Alexa Fluor 594, goat anti-chicken Alexa Fluor 488 and goat anti-mouse Alexa Fluor 568 (1:250; Invitrogen, Thermo Fisher Scientific). Sections were washed and further incubated with 4’,6-diamidino-2-phenylindole dihydrochloride (DAPI; 1:5000; Sigma-Aldrich), before being placed in gelatine-coated slides and coverslipped with fluorescence mounting medium (Dako, Agilent Technologies, Santa Clara, CA, USA).

Images were captured with a slide scanner Zeiss Axio Scan.Z1 microscope (Carl Zeiss Microscopy GmbH), equipped with a Hamamatsu ORCA-Flash 4.0 camera, using the Plan-Apochromat 20x/0.8 objective and the ZEN software (Carl Zeiss Microscopy GmbH).

### Immunohistochemical analysis: Quantification of ubiquitin-positive inclusions and measurement of DARPP-32-depleted volume (lentiviral-based model)

Twelve stained sections per animal, distanced by 200 µm from each other, were used to perform the quantitative analysis of the total number of ubiquitin-positive inclusions as well as the extent of DARPP-32 loss in the striatum. Total number of inclusions was manually counted in all 12 sections and further multiplied by 8 to account for the intermediate sections. The quantification of DARPP-32 depleted areas in mice striatal sections was performed with an image-analysis software (ImageJ software, National Institutes of Health, Bethesda, MD, USA). The calculation of DARPP-32-depleted volume was based on the following formula: 𝑉𝑜𝑙𝑢𝑚𝑒 = 𝑑(𝑎1 + 𝑎2 + ⋯ + 𝑎12), where *d* is the distance between serial sections and *a1* to *a12* represent the individual areas for serial sections.

### Immunohistochemical analysis: Quantification of Iba-1 and Gfap immunoreactivity (lentiviral-based model)

Quantitative analysis of the Iba-1 and Gfap immunoreactivity was performed with ZEN 2 Blue edition software (Carl Zeiss Microscopy GmbH) in the transduced striatal regions relative to their corresponding non-affected cortex (defined as background). Twelve stained-sections per animal, distanced by 200 µm from each other, were used for the purpose. Images were taken under the same image acquisition conditions and uniform adjustments of brightness and contrast were made to all images.

### Immunohistochemical analysis: Quantification of ATXN3 immunoreactivity (YAC-MJD84.2/84.2 mice)

Nuclear ATXN3 accumulation was quantified in the DCN region over a total of seven or four (adult or neonatal YAC-MJD84.2/84.2-injected mice, respectively) sagittal sections (right hemispheres). Image quantification was performed using the QuPath software version 0.5.1 ^84^, in a semi-automated fashion. The total number of cells with ATXN3-positive inclusions (“cell detections”) was further multiplied by 8 to account for the intermediate sections. Annotations corresponding to the deep cerebellar nuclei were manually delineated. Images were taken under the same image acquisition conditions and uniform adjustments of brightness and contrast were made to all images.

### Statistical analysis

One-way analysis of variance (ANOVA) followed by Dunnett’s *post hoc* test was used to compare protein levels in the human cell line transfected with TALENs and CRISPR systems. Paired Student’s t-test was used in the lentiviral-based models. Statistical comparisons of YAC-MJD84.2/84.2 mice body weight were performed using a mixed effects analysis followed by the adequate *post hoc* test for multiple comparisons. Motor performance of YAC-MJD84.2/84.2 mice, revealed on the open-field test, was analyzed using either a two-way ANOVA with post hoc comparisons by the Tukey test (YAC-MJD84.2/84.2 neonatal mice, ICM-injected) or the mixed effects analysis followed by Tukey’s *post hoc* test (YAC-MJD84.2/84.2 mice injected in the cerebellar parenchyma). The number of viral genomes in the different brain regions and the number of cells with ATXN3 nuclear accumulation (YAC-MJD84.2/84.2 neonatal mice, ICM-injected) were analyzed using unpaired Student’s t-test. For the remaining analysis, statistics was performed using the one-way ANOVA followed by Tukey’s multiple comparison test. Data are presented as mean ± standard error of mean (SEM). Significance thresholds were set at *p<0.05, **p<0.01, ***p<0.001 and ****p<0.0001 and considered non-significant (ns) when p>0.05. Calculations were performed using GraphPad Prism version 8.4.0 for Windows (San Diego, CA, USA).

