## Supplemental Figures and Tables for "Gene Editing for *ATXN3* Inactivation in Machado-Joseph disease: CRISPR-Cas9 as a Therapeutic Alternative to TALEN-Induced Toxicity"

### SUPPLEMENTAL FIGURES

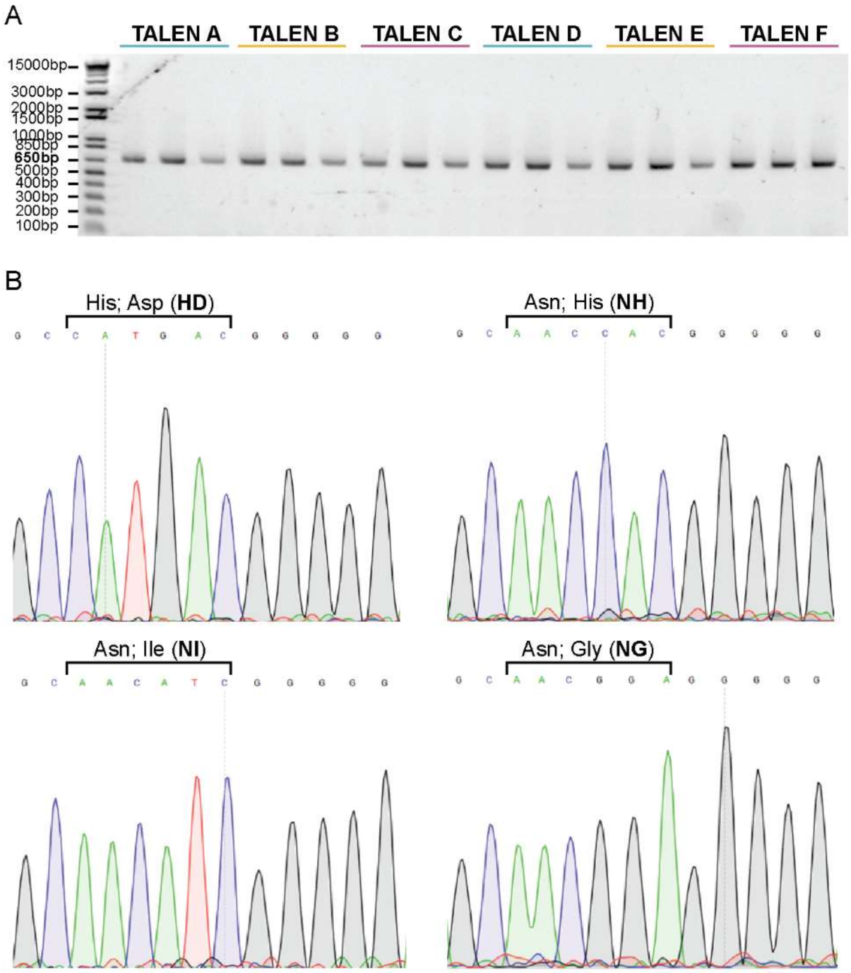


**Supplemental Figure S1 – Checkpoints for correct TALEN assembly (A)** Agarose gel showing the presence of 650 bp bands, corresponding to PCR-amplified hexamers, successfully assembled by Golden Gate cut-ligation. Lane 1: 1-kb Plus DNA ladder; Lane 2-4: hexamers for TALEN A assembly; Lane 5-7: hexamers for TALEN B assembly; Lane 8-10: hexamers for TALEN C assembly; Lane 11-13: hexamers for TALEN D assembly; Lane 14-16: hexamers for TALEN E assembly; Lane 17-19: hexamers for TALEN F assembly; **(B)** Representative electropherograms displaying residues at positions 12 and 13 (RVDs) of each monomer for the verification of the correct assembly of each TALEN.

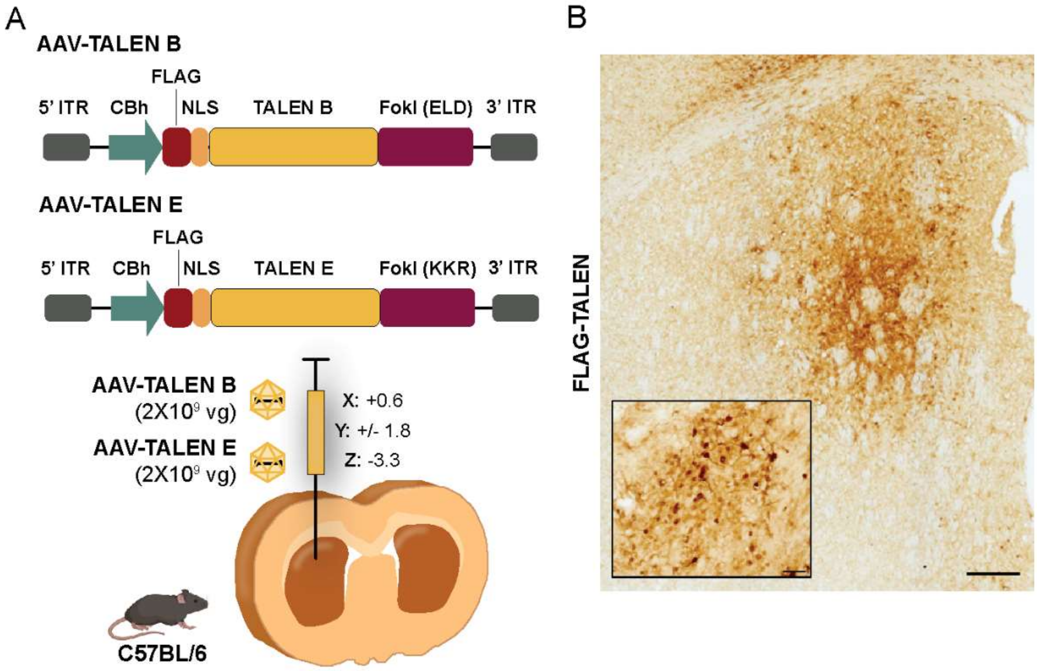


**Supplemental Figure S2 – rAAV1/2 vectors encoding TALENs B and E efficiently transduce the mouse striatum. (A)** Graphical representation of the stereotaxic procedure. rAAVs encoding TALENs B and E were injected into the striatum of seven-week-old C57BL/6 mice at the following coordinates (X: + 0.6; Y: +/- 1.8; Z: -3.3). These vectors were efficient to mediate *in vivo* transduction of the brain parenchyma 4 weeks after injection, as can be observed through the **(B)** positive staining of the FLAG-tagged TALEN constructs. Scale bar, 200 µm and 50 µm in the largest and smallest images, respectively.

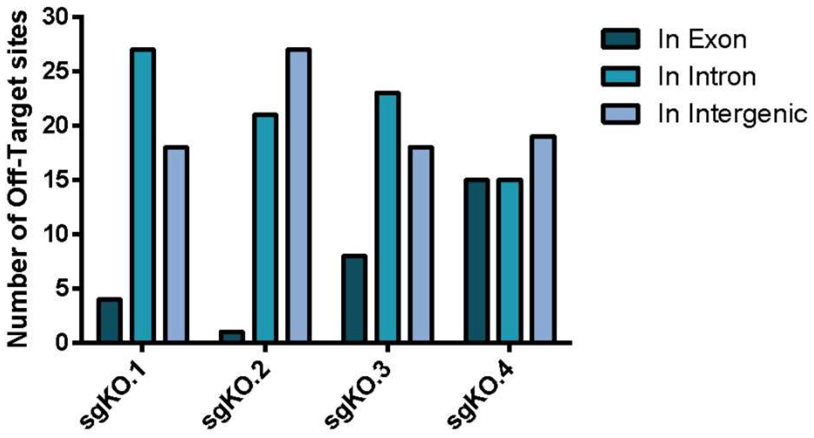


**Supplemental Figure S3 – Bioinformatic analysis of potential off-targets in the human genome for the sgKO sequences.** Fifty potential off-target sites, scored in accordance with the likelihood of off-target binding, were predicted for each guide sequence, using the Benchling bioinformatic tool. Genomic locations of the predicted off-targets were determined using the UCSC genome browser and mapped to intergenic, intronic and exonic regions. The large majority of putative off-targets are located in intergenic or intronic regions.

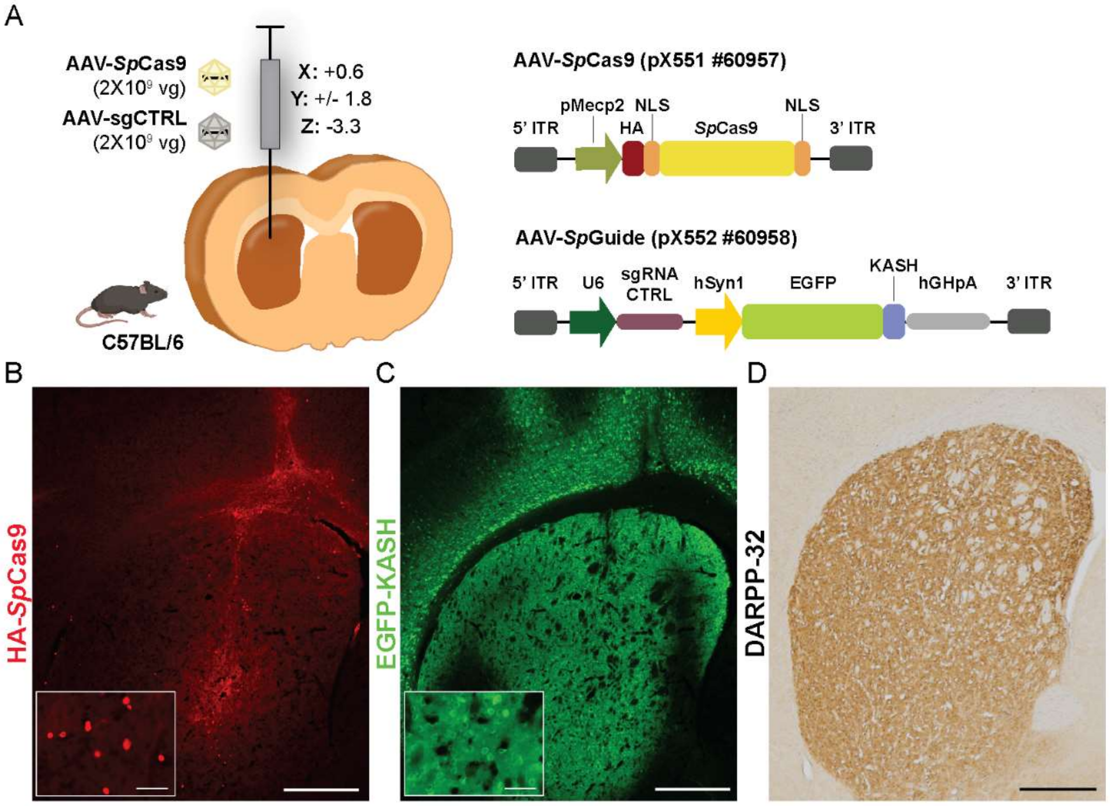


**Supplemental Figure S4 – Co-delivery of rAAV1/2-*Sp*Cas9 and rAAV1/2-sgCTRL *in vivo* effectively transduces the mouse brain with no associated toxicity. (A)** rAAVs encoding for *Sp*Cas9 (pX551, addgene plasmid #60957) and the control guide sequence (cloned in *Sp*Guide, pX552, addgene plasmid #60958) were injected in the striatum of seven-week-old C57BL/6 mice at the following coordinates (X: +0.6; Y: +/-1.8; Z: -3.3). These vectors were efficient to mediate *in vivo* transduction of the brain parenchyma 4 weeks after injection, as can be observed through the expression of **(B)** HA-tagged *Sp*Cas9 and of **(C)** EGFP-KASH fusion protein, encoded in the *Sp*Guide plasmid. **(D)** No loss in DARPP-32 immunoreactivity was observed in the injected striatal hemisphere, suggesting no rAAV1/2-associated neurotoxicity (n=2). Scale bar, 500 μm in general views and 50 μm in detail magnifications.

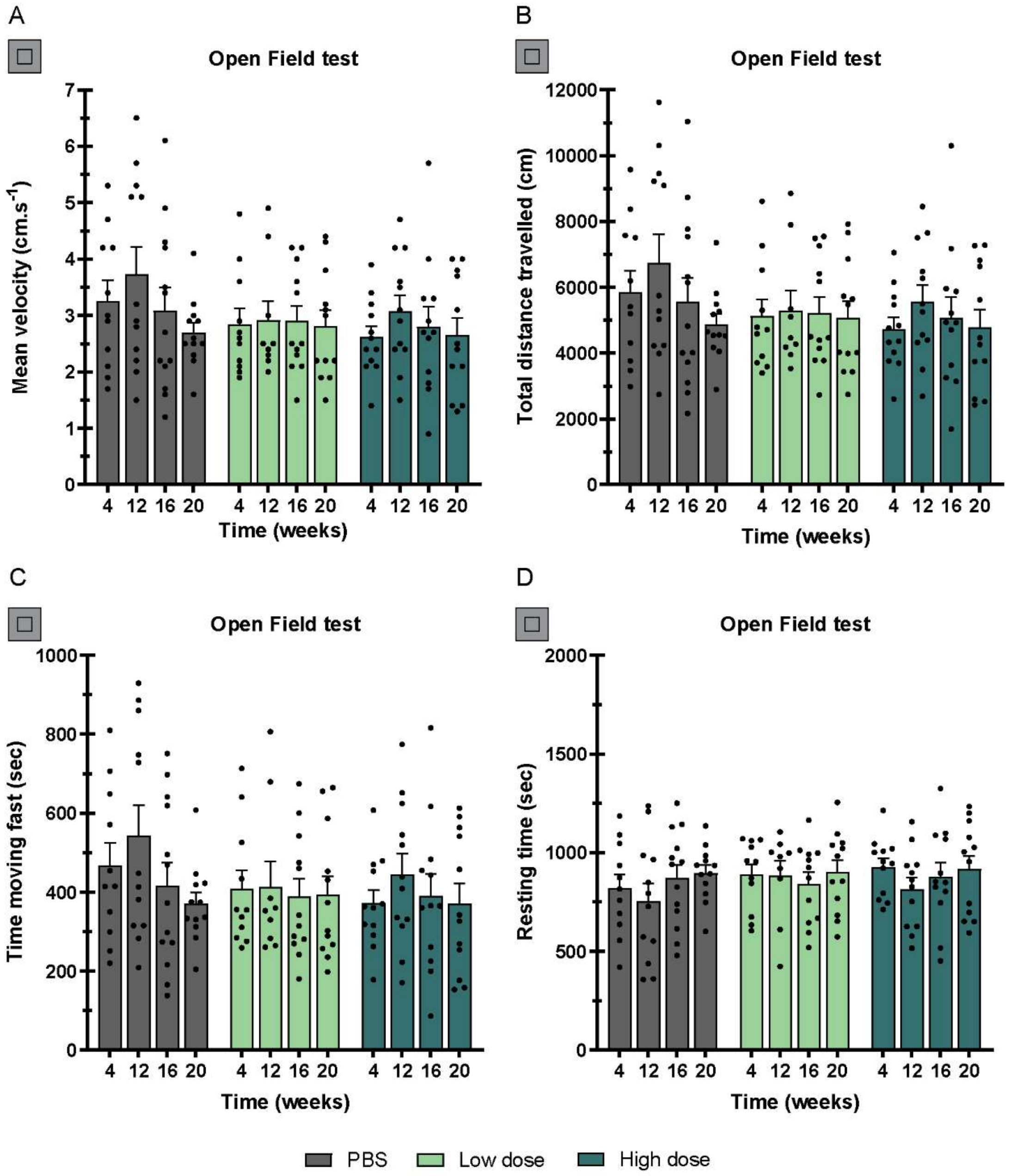


**Supplemental Figure S5 –Post-symptomatic administration of the CRISPR-*ATXN3* KO system into the cerebellum of adult YAC-MJD mice did not improve motor deficits in the open-field test.** Locomotor horizontal activity over a 30-minute trial in the open field test. No statistically significant differences are observed in-between the groups of animals, in any of the assessed parameters: **(A)** mean velocity (in $cm.s^{-1}$), **(B)** total distance travelled (in centimeters), **(C)** time (in seconds) moving fast (above 5 $cm.s^{-1}$), and **(D)** resting time (in seconds). Values are presented as mean ± SEM. Mixed effects analysis with post-hoc Tukey's multiple comparison test was performed (n=13 for the control group and n=12 for the treatment groups).

### SUPPLEMENTAL TABLES

### Supplemental Table 1 – TALEN target sequences divided into hexamers

| | | Target Sequence (5'-$T_0N_{19}$-3') | Hexamer Modules | |
|---|---|---|---|---|
| Left TALENs | TALEN A | TTAGCAAGAAGGCTCACTTT | $N_1N_2N_3N_4N_5N_6$ | NG.NI.NH.HD.NI.NI |
| | | | $N_7N_8N_9N_{10}N_{11}N_{12}$ | NH.NI.NI.NH.NH.HD |
| | | | $N_{13}N_{14}N_{15}N_{16}N_{17}N_{18}$ | NG.HD.NI.HD.NG.NG |
| | | | $N_{19}$ (0.5 repeat) | NG |
| | TALEN B | TATTGCAAGGAGAATATTTT | $N_1N_2N_3N_4N_5N_6$ | NI.NG.NG.NH.HD.NI |
| | | | $N_7N_8N_9N_{10}N_{11}N_{12}$ | NI.NH.NH.NI.NH.NI |
| | | | $N_{13}N_{14}N_{15}N_{16}N_{17}N_{18}$ | NI.NG.NI.NG.NG.NG |
| | | | $N_{19}$ (0.5 repeat) | NG |
| | TALEN C | TGGAAATATTAGCAAGAAGG | $N_1N_2N_3N_4N_5N_6$ | NH.NH.NI.NI.NI.NG |
| | | | $N_7N_8N_9N_{10}N_{11}N_{12}$ | NI.NG.NG.NI.NH.HD |
| | | | $N_{13}N_{14}N_{15}N_{16}N_{17}N_{18}$ | NI.NI.NH.NI.NI.NH |
| | | | $N_{19}$ (0.5 repeat) | NH |
| Right TALENs | TALEN D | TCCTTGCAATAAGTTATTCA | $N_1N_2N_3N_4N_5N_6$ | HD.HD.NG.NG.NH.HD |
| | | | $N_7N_8N_9N_{10}N_{11}N_{12}$ | NI.NI.NG.NI.NI.NH |
| | | | $N_{13}N_{14}N_{15}N_{16}N_{17}N_{18}$ | NG.NG.NI.NG.NG.HD |
| | | | $N_{19}$ (0.5 repeat) | NI |
| | TALEN E | TCCAGCTGATGTGCAATTGA | $N_1N_2N_3N_4N_5N_6$ | HD.HD.NI.NH.HD.NG |
| | | | $N_7N_8N_9N_{10}N_{11}N_{12}$ | NH.NI.NG.NH.NG.NH |
| | | | $N_{13}N_{14}N_{15}N_{16}N_{17}N_{18}$ | HD.NI.NI.NG.NG.NH |
| | | | $N_{19}$ (0.5 repeat) | NI |
| | TALEN F | TGCAATAAGTTATTCAGGCA | $N_1N_2N_3N_4N_5N_6$ | NH.HD.NI.NI.NG.NI |
| | | | $N_7N_8N_9N_{10}N_{11}N_{12}$ | NI.NH.NG.NG.NI.NG |
| | | | $N_{13}N_{14}N_{15}N_{16}N_{17}N_{18}$ | NG.HD.NI.NH.NH.HD |
| | | | $N_{19}$ (0.5 repeat) | **NI** |

### Supplemental Table 2 – Top and bottom oligonucleotides encoding sgRNA sequences

| Plasmid | Target Locus | Oligo's nomenclature | Oligo's Sequence (5' → 3') |
|---|---|---|---|
| lentiCRISPRv2 (#52961) | *ATXN3* gene (Exon 2) | sgKO.1 top | CAC CGA TCC TCA ATT GCA CAT CAG C |
| | | sgKO.1 bottom | AAA CGC TGA TGT GCA ATT GAG GAT C |
| | | sgKO.2 top | CAC CGT GCC TGA ATA ACT TAT TGC A |
| | | sgKO.2 bottom | AAA CTG CAA TAA GTT ATT CAG GCA C |
| | | sgKO.3 top | CAC CGA ATT GCA CAT CAG CTG GAT G |
| | | sgKO.3 bottom | AAA CCA TCC AGC TGA TGT GCA ATT C |
| | | sgKO.4 top | CAC CGA TCA GCT GGA TGA GGA GGA G |
| | | sgKO.4 bottom | AAA CCT CCT CCT CAT CCA GCT GAT C |
| | Bacterial *LacZ* gene (Non-Targeting Control guide) | sgCTRL top | CAC CGT GCG AAT ACG CCC ACG CGA T |
| | | sgCTRL bottom | AAA CAT CGC GTG GGC GTA TTC GCA C |
| AAV - *SpGuide* (pX552 #60958) | *ATXN3* gene (Exon 2) | sgKO.2 top | ACC GTG CCT GAA TAA CTT ATT GCA |
| | | sgKO.2 bottom | AAC TGC AAT AAG TTA TTC AGG CAC |
| | Bacterial *LacZ* gene (Non-Targeting Control guide) | sgCTRL top | ACC GTG CGA ATA CGC CCA CGC GAT |
| | | sgCTRL bottom | AAC ATC GCG TGG GCG TAT TCG CAC |

**Supplemental Table 3 – Expected sizes for the Surveyor nuclease-cleaved products**

| Target Gene | PCR amplicon (bp) | Surveyor nuclease digestion products (bp) |
|---|---|---|
| ***ATXN3*** (genomic) | 762 | **TALEN pair A/D:** ~ 92 + 670 |
| | | **TALEN pair B/E:** ~ 138 + 624 |
| | | **TALEN pair C/F:** ~ 86 + 676 |
| | | **sgKO.1:** 160 + 602 |
| | | **sgKO.2:** 113 + 649 |
| | | **sgKO.3:** 166 + 596 |
| | | **sgKO.4:** 174 + 588 |
| **LV-PGK-*ATXN3* 72Q** (cDNA) | 485 | **TALEN pair B/E:** ~ 163 + 322 |
| | | **sgKO.2:** 137 + 348 |
